# Knockdown of the long isoform of the prolactin receptor selectively targets pathogenic immune cells in systemic lupus erythematosus and averts glomerular pathology

**DOI:** 10.64898/2026.03.12.711422

**Authors:** Kory Hamane, Zunsong Hu, Joao Rodrigues Lima-Junior, Adeleh Taghi Khani, Anil Kumar, Ashly Sanchez Ortiz, Xinying Guo, Da Hae Jung, Hanjun Qin, Shu Tao, Ignacio Sanz, Eric Meffre, Katherine Marzan, Jean L. Koff, Mary Y. Lorenson, Zhaohui Gu, Xiwei Wu, Ameae M. Walker, Srividya Swaminathan

## Abstract

Systemic lupus erythematosus (SLE) is an autoimmune disease characterized by chronic inflammation in multiple organ systems. While a clinical association between elevated levels of the hormone/cytokine, prolactin (PRL), and exacerbation of SLE has been recognized for some time, little is known about the mechanisms through which PRL affects the course of this disease. Here, we show that immune cells in SLE have aberrant splicing of the prolactin receptor (PRLR) such that the ratio of the long to short splice variants is increased. To determine whether the change in PRLR isoform expression was causal in this disease, we used a splice-modulating oligomer (SMO) that knocked down expression of the long splice variant (LFPRLR). Using patient samples *ex vivo*, SLE-prone mice *in vivo*, high-dimensional flow cytometry and single-cell RNA-sequencing, we demonstrate that the aberrant PRLR isoform expression in SLE both directly and indirectly drives production of autoreactive immune cell phenotypes. Thus, LFPRLR knockdown decreased the expression of type I interferon signaling/response genes known to be biomarkers and hub genes for SLE, reduced immunoglobulins with signatures considered autoreactive, and averted glomerular kidney damage in SLE-prone mice. Importantly, LFPRLR knockdown reduced pathogenic B cells and other immune subsets that drive B-cell activation, without negative impact on healthy donor counterparts. Current treatments for SLE adversely affect healthy cells and do not concurrently eradicate multiple pathogenic immune subsets. Since LFPRLR SMO does not share these disadvantages, knockdown of the LFPRLR represents a potential treatment strategy for SLE that merits further investigation.

**Graphical Abstract:** **The LFPRLR represents an attractive therapeutic target in SLE.** (**Left**) In addition to increased pituitary/circulating PRL, individuals with SLE exhibit aberrant increases in the production of autocrine/paracrine PRL by immune cells, and in their expression of specifically the long isoform (LF) of the PRLR. (**Middle**) Expression of the LFPRLR specifically enhances autoreactive immunophenotypes and promotes lupus nephritis. (**Right**) A splice modulating oligomer (SMO), that prevents synthesis of only the LFPRLR but not the short PRLR isoforms, reduces pathogenic immunophenotypes without affecting the normal counterparts of immune cells.

## Introduction

Systemic lupus erythematosus (SLE) is an autoimmune disease in which chronic activation of the immune system induces inflammation in multiple tissues, including the kidneys, skin, joints, heart, lungs, and brain^1^. Tissue inflammation and malfunction are triggered by abnormal B cells and their autoantibody-secreting derivatives, plasma cells^2^. Autoreactive B cells do not function in isolation; both their production and pathogenic effects are mediated by other inflammatory immune cells^1^.

Current treatments, including anti-inflammatory drugs, immunosuppressive steroids, pan-B-cell-depleting regimens, and plasma cell-depleting agents, do not sustain SLE remission because they are neither specific to pathogenic cells nor do they concurrently target multiple pathogenic immune subsets^1^. For example, pan-B-cell-depleting agents do not eradicate autoantibody-secreting plasma cells^2,3^. They are therefore alternated with agents that only eradicate plasma cells^3^. As a result, patients with SLE receive multiple therapies and experience a variety of morbidities resulting from their side effects. Understanding more about what drives this disease may lead to better therapies.

Increased circulating prolactin (PRL) is positively correlated with worsening SLE symptoms, and administration of the dopamine agonist, bromocriptine, can improve patient symptomatology^4,5^. Circulating PRL levels are higher in females than males under most circumstances, in large part because estrogen increases pituitary PRL secretion ^reviewed^ ^in^ ^6^. Since dopamine agonists do not affect estrogen levels, the improvement of symptomatology with bromocriptine is the result of effects on levels of PRL and not estrogen^4,5^. Higher PRL levels correlate with the higher incidence of SLE in females compared to males^7^. However, estrogen is not the only regulator of PRL release; both psychological and physical stressors, for example, elevate circulating PRL in both males and females^6^. Thus, PRL can be a major contributor to the pathogenesis of SLE in both sexes^6^.

Most PRL is produced in the pituitary, but many other tissues, including some immune cells and immune tissue stromal cells, produce smaller quantities^6^. Dopamine agonists, which limit secretion of pituitary PRL and, as mentioned above ameliorate symptoms of SLE in some patients^4,5^, do not affect the release of extrapituitary PRL^8^. Thus, to interfere with the action of all PRL, the prolactin receptor (PRLR) represents a better target.

PRL-responding cells express alternatively spliced PRLRs *viz*. a long (LF), intermediate (IF, only described in pathological specimens), and two to three short (SF) isoforms^9,10^. Each of these PRLR isoforms includes exons 3-9, which code for identical extracellular and transmembrane domains. Exon 10 codes for a portion of the receptor essentially only present in LF/IF mRNA, while splicing to produce the SFs attaches different portions of exon 11 (human) or exons 11-13 (mouse) to exon 9 to result in the intracellular signaling domains^11^. Thus, PRL signaling can produce different outcomes depending on the isoforms of PRLR present. Work from multiple laboratories has shown that increased relative expression of the LF/IF leads to cell proliferation and survival, whereas increased relative expression of one or more of the SFs decreases proliferation and increases differentiation and apoptosis^6^.

Consistent with the positive association between elevated levels of circulating PRL and exacerbation of symptoms in patients with SLE^6^, administration of exogenous PRL or PRL secretagogues was found to induce some B-cell and other pathogenic immune subsets in mouse models of SLE^12,13^. However, how PRL drives autoreactive processes was unknown. Here we have characterized the PRLR isoforms in each immune subset and, in so doing, have demonstrated aberrant splicing of the PRLR in SLE, producing a higher LF to SF PRLR ratio in diseased cells.

Further, use of a non-toxic, patented, splice-modulating oligomer (SMO)^14,15^ that decreases the LF:SF PRLR ratio (termed LFPRLR SMO) demonstrated that aberrant PRLR splicing was causal in this disease; treatment with the LFPRLR SMO markedly reduced pathogenic immune subsets and averted development of glomerular pathology.

## Results

### Immune cells in SLE have aberrant splicing of PRLRs with an increase in the ratio of the long to short splice variants

Bulk RNA-sequencing (RNA-seq) analysis of PBMCs^16^ from SLE patients (n=20) and healthy donors (n=10) revealed an increase in total PRLR expression, specifically the LFPRLR (identified by the presence of exon 10), in SLE (**Fig. 1A left and middle**). In contrast, PRL expression in total immune cells was not different (**Fig. 1A right**). Analysis (outlined in **Fig. S1**) by single-cell RNA-sequencing (scRNA-seq)^16^ of healthy donors (n=12) and SLE patient (n=34) PBMCs demonstrated that the increase in PRLR in SLE total PBMCs is driven more by elevated expression levels within PRLR^+^ immune cells than by an expansion in the proportion of PRLR^+^ cells (**Fig. 1B**), i.e., particular cells increase their expression in SLE rather than there being more cells expressing the receptor. Consistent with bulk RNA-seq data (**Fig. 1A right panel**), no overall change in PRL expression in SLE immune cells compared to their normal counterparts was seen in scRNA-seq data (**Fig. 1C left**) likely because the reduction in proportion of cells expressing PRL (**Fig. 1C middle**) was balanced by an increase in PRL expression within individual PRL^+^ immune cells in SLE (**Fig. 1C right**).

**Fig 1.**
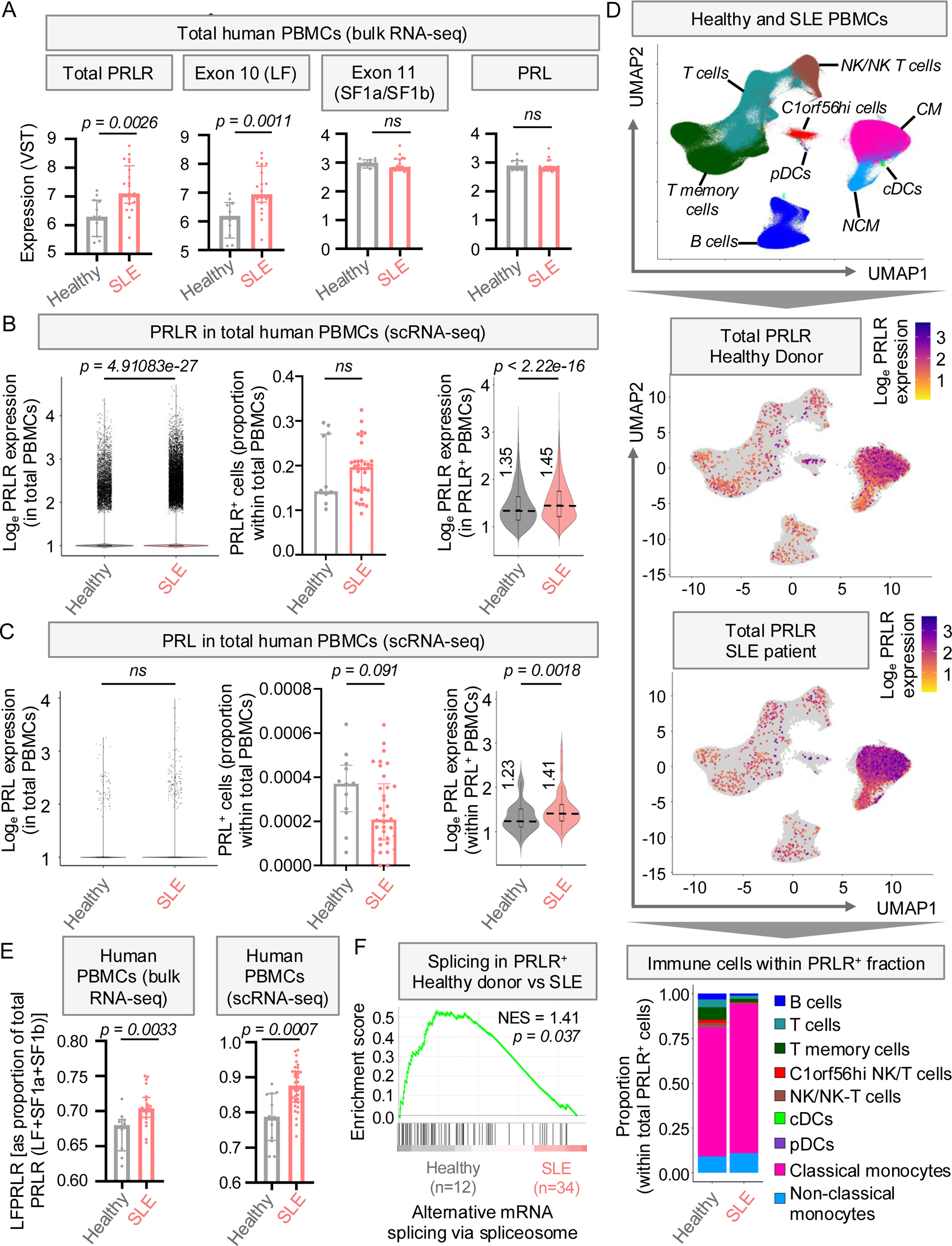
Immune cells in SLE have aberrant splicing of PRLRs with an increase in the ratio of the long to short splice variants. (**A**) Total PRLR, LFPRLR, SFPRLR, and PRL expression in PBMCs from human patients with SLE (n=20) and healthy donors (n=10) by bulk RNA-seq (GSE211700). Median ± interquartile range, p-values: Mann-Whitney U test. (**B-C**) scRNA-seq (GSE137029) depicting overall distribution of expression of PRLR and PRL transcripts (left), proportion of PRLR^+^ or PRL^+^ cells (middle), and expression of PRLR and PRL transcripts within PRLR^+^ and PRL^+^ cells (right) in PBMCs of healthy donors (n=12) and patients with SLE (n=34). Left panel p-value: Seurat’s implementation of the likelihood ratio test for bimodal distribution; right and middle panel p-values: Mann-Whitney U test. (**D**) UMAP dimensional reduction of 524,031 PBMCs from 34 patients with SLE and 209,597 PBMCs from 12 healthy donors showing immune cell types, expression of PRLR, and the distribution of immune cell types in PRLR^+^ PBMCs by scRNA-seq. (**E**) Bulk RNA-seq and scRNA-seq depicting the relative LFPRLR (as a proportion of total PRLR [LF+SF1a+SF1b]) in PBMCs from healthy donors (n= 10 for bulk RNA-seq, n= 12 for scRNA-seq) and patients with SLE (n=20 for bulk RNA-seq, n=34 for scRNA-seq). Median ± interquartile range, p-values: Mann-Whitney U test. (**F**) GSEA comparing the pathways/gene ontology gene set of ‘alternative mRNA splicing via the spliceosome’ in PRLR^+^ PBMCs of healthy donors (n=12) and patients with SLE (n=34). P-values were calculated using the permutation test within GSEA.

In both healthy and SLE samples, human monocytes, particularly the classical monocyte (CM) fraction, express the most PRLR (**Fig. 1D**), which is predominantly the LF (**Fig. S2A**). The proportion of PRLR^+^ monocytes was significantly expanded at the expense of PRLR^+^ lymphoid subsets in SLE patients (**Figs. 1D, S2B**). There is therefore an increased potential for monocytes to respond to PRL in SLE.

Both bulk and scRNA-seq data of healthy donor and SLE patient PBMCs showed that the balance of the PRLRs is tilted significantly in favor of LF in SLE (**Fig. 1E**). Gene set enrichment analysis (GSEA) revealed that expression of transcripts involved in alternative splicing was reduced in PRLR^+^ but not in PRLR^−^ SLE PBMCs compared to their healthy counterparts (**Fig. 1F**). The unusual alteration of the spliceosome only in PRLR^+^ immune cells is concordant with the skewing towards LFPRLR rather than SFPRLR production in SLE.

Consistent with what was found in humans, bulk RNA-seq of tissues from *NZB/W-F1* SLE-prone mice^17^ revealed that LFPRLR was predominant in the spleen and increased with disease progression/age in splenocytes compared to controls (**Figs. S3A-C**). The increase in LFPRLR did not occur in the brain or kidney of SLE-prone mice (**Figs. S3A-C**). These data suggest at least a somewhat immune-focused change in PRLR signaling in SLE and provide a rationale for interrogating the cause-and-effect relationship between PRLR isoform changes and immune cell pathology in this disease.

### *Ex vivo* knockdown of the LFPRLR in PBMCs from patients with SLE reduces pathogenic immunophenotypes but has no negative impact on healthy PBMCs

To determine whether the change in PRLR splice variants was causal in disease, we treated PBMCs from patients with SLE and age-and sex-matched cells from healthy donors with the LFPRLR SMO *ex vivo*. The LFPRLR SMO, is a morpholino DNA oligonucleotide derivatized with octaguanidine to promote cell uptake^13^. We have previously demonstrated that the LFPRLR SMO changes the PRLR isoform expression by reducing the LF and/or increasing the SFs in both cancer models and in mice prone to SLE^14,15^.

In the present study, we first confirmed the functionality of LFPRLR SMO by treating healthy donor (HD) and SLE patient PBMCs and analyzing receptor variant expression by exon junction qPCR, which distinguishes between the LF and SFs. Consistent with its on-target splice modulation, LFPRLR SMO treatment reduced the proportion of LF within the total PRLR in both healthy donor and SLE patient PBMCs compared to their control SMO-treated counterparts (**Figs. 2A, S4**). Then, using high-dimensional flow cytometry (gating strategy in **Fig. S5**), we compared the effects of a 3-day *ex vivo* knockdown of the LFPRLR in PBMCs of SLE patients (n=9, patient characteristics in **Table S1**) and age-and sex-matched healthy donors (n=8). Two of the SLE patients had received short-term (<2 months), and seven patients had received long-term (≥ 2 months) immunomodulatory/immunosuppressive non-steroid and/or steroid regimens. Treatment-naïve samples were unavailable. The effect of LFPRLR SMO on immune subsets in each sample was computed as a ratio of the frequency of the immune subset and/or its expression of a given marker in LFPRLR SMO *vs* control SMO group. The derived ratios for both healthy and SLE samples were compared (**Fig. 2B**). This computation ensures that the effects of LFPRLR knockdown are not confounded by the heterogeneity in disease activity [as measured by the SLE Disease Activity Index (SLEDAI) score], treatment type and duration, and age of the patient.

**Fig 2.**
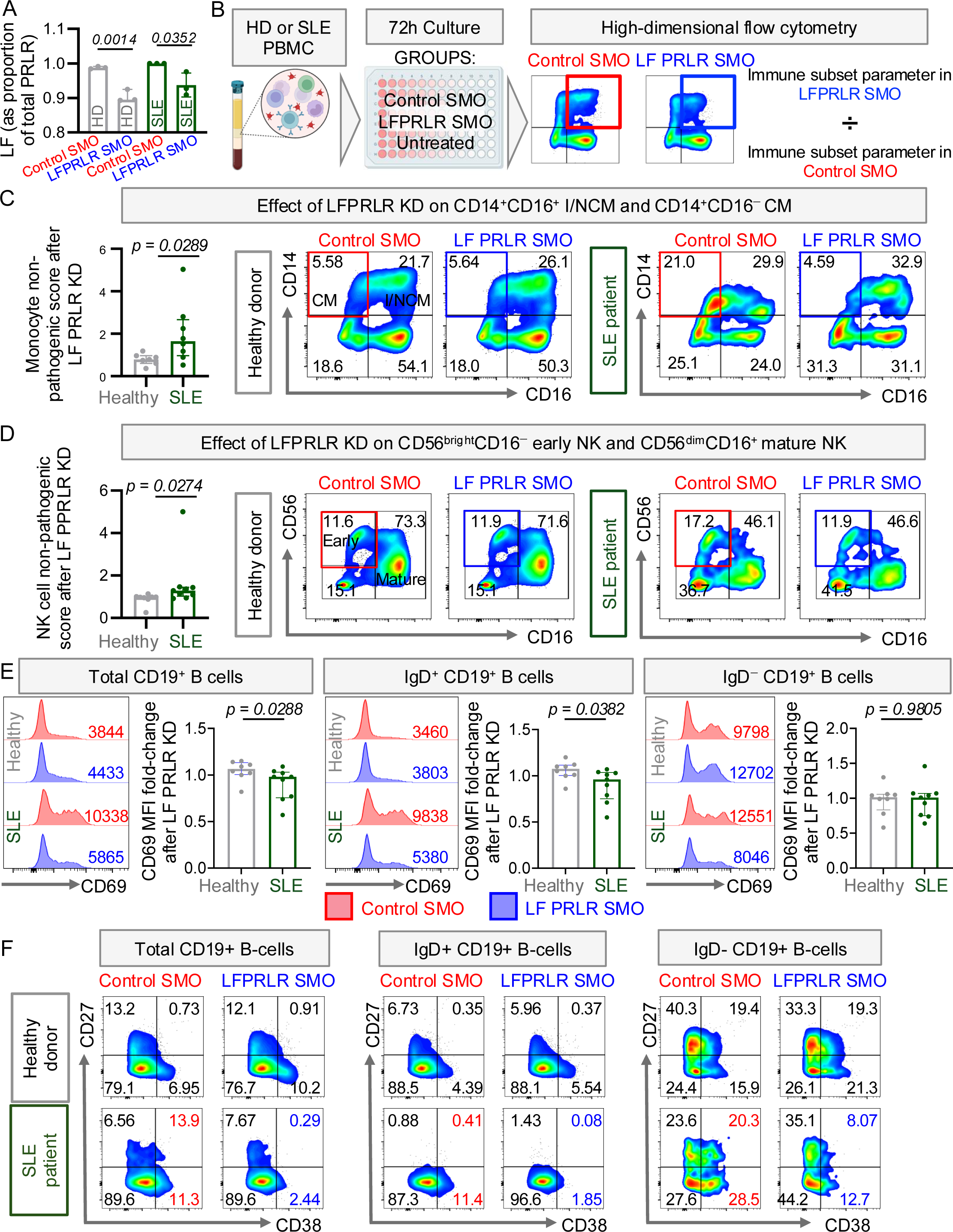
*Ex vivo* knockdown of the LFPRLR in PBMCs from SLE patients reduces pathogenic immunophenotypes but has no negative impact on healthy PBMCs. (**A**) Relative LFPRLR (as a fraction of total PRLR) was quantified by qPCR in healthy donor (HD) and SLE patient PBMCs treated with control SMO or LFPRLR SMO for 24h. Transcripts were normalized to ubiquitin (UBB) and expressed as % of UBB. Each dot represents a technical replicate within a single healthy donor (grey) or SLE patient (green). One representative of three independent experiments, each containing one HD and one age-and sex-matched SLE sample, is shown. Median ± interquartile range, p-values: unpaired two-tailed Student’s t test. ns = not significant. (**B**) Experimental workflow for measuring the effects of *ex vivo* knockdown of LFPRLR for 72h in PBMCs of healthy donors and patients with SLE by high-dimensional flow cytometry. (**C-D**) Effect of LFPRLR knockdown on the ratio of (**C**) less mature classical monocytes (CMs, (CD14⁺CD16⁻) to more mature intermediate and non-classical monocytes (I/NCM, CD14⁺CD16⁺) monocyte subsets in PBMCs from healthy donors (n = 7) and patients with SLE (n=8), and (**D**) immature (CD56ᵇʳⁱᵍʰᵗCD16⁻) to more-mature (CD56ᵈⁱᵐCD16⁺) NK-cell subsets in PBMCs from healthy donors (n=9) and patients with SLE (n=9). For (**C-D**), the effect of LF PRLR KD on monocyte and NK cell non-pathogenic phenotype score was calculated as: (% of more mature ÷ % of less mature subset in LF PRLR SMO-treated sample) ÷ (% of more mature ÷ % of less mature subset in control SMO-treated sample). (**E**) Fold-change in CD69 median fluorescence intensity (MFI) in total, IgD⁺, and IgD⁻ CD19⁺ B cells from healthy donors (n=8) and patients with SLE (n=9), calculated as the ratio of MFI in LFPRLR SMO-treated cells to that in control SMO-treated cells. (**F**) Percentages of CD27/CD38 subsets within total, IgD⁺, and IgD⁻ CD19⁺ B cell fractions in healthy donors (n=7) and patients with SLE (n=8) after treatment with control or LFPRLR SMO. Median ± interquartile range, p-values: Mann-Whitney U test.

After confirming that *ex vivo* knockdown of the LFPRLR for 72 hours did not affect the viability of either healthy donor or SLE patient PBMCs (**Fig. S6**), we examined the effects of knockdown on monocytes because their PRLR expression is highest among the cell types in human PBMCs (**Figs. 1D, S2A**) plus monocytes can trigger autoantibody production^18,19^. Looking at healthy donor cells, one can appreciate that the LFPRLR SMO had no effect (**Figs. 2C**). However in the SLE samples, the proportion of more mature, CD14^+^CD16^+^, intermediate/non-classical monocytes (I/NCM) to the less mature, CD14^+^CD16^−^ CM^20,21^ was markedly increased after LFPRLR knockdown (**Fig. 2C**).

Natural killer (NK) cells that are CD56^bright^CD16^−^ (termed ‘early NK’ in **Fig. 2D**) are less mature and poorly cytotoxic but produce more inflammatory cytokines compared to their CD56^dim^CD16^+^ counterparts (termed ‘mature NK’ in **Fig. 2D**). CD56^bright^CD16^−^ NK cells are increased in response to type I interferons (IFN-Is) in patients with SLE^22^, further enhance IFN-I production by plasmacytoid dendritic cells (pDCs)^23^, and stimulate B-cell activation in SLE^24,25^. In contrast, CD56^+^CD16^+^ NK cells eradicate autoreactive B cells^26,27^. We found that the population of NK cells is more mature in healthy donors *versus* SLE patients and that LFPRLR knockdown once again had no effect on healthy donor cells (**Fig. 2D**). As anticipated, SLE patients had more CD56^bright^CD16^−^, and treatment with LFPRLR SMO for 72 hours reduced the proportion of immature cells to levels seen in healthy donors (**Fig. 2D**).

Inflammatory monocytes and NK cells drive the activation and generation of self-reactive B cells. Thus, non-class-switched IgD^+^ B cells increase expression of CD69, become activated, and mature into class-switched memory B cells and plasma cells^28–30^. Higher surface CD69 expression was found in total and CD19^+^ B-cell subsets in SLE. LFPRLR knockdown significantly reduced the expression of CD69 on total and non-class switched IgD^+^ B cells in SLE patient samples, without a significant effect on the healthy donor counterparts (**Fig. 2E**). Thus, LFPRLR knockdown reduces aberrant B-cell activation and therefore could block the subsequent generation of autoantibody-producing plasma cells.

Some changes did not reach statistical significance when considering all patients together, likely the result of variation among patients in SLEDAI score^31^, treatment type, and duration of treatment (**Table S1**). For example, LFPRLR knockdown reduced the proportion of IgD^−^CD19^+^ B cells expressing CD38 in three SLE patients without impact on the corresponding age-matched healthy donors (**Fig. 2F**). In one sample from a patient who presented multiple autoimmune pathologies, including SLE, Sjogren’s Syndrome, hypothyroidism, and Raynaud’s syndrome (Patient 23150, **Table S1**), LFPRLR knockdown reduced several other pathogenic immune populations in addition to those shown in **Fig. 2**. These included reductions in a) classical IFN-I-producing pDCs and conventional dendritic cells (cDCs), which also produce some IFN-Is (**Fig. 3A**); b) CD137^+^CXCR3^+^ atypical B cells^32,33^ (**Fig. 3B**); and c) CD38^+^CD138^+/−^ B cells (**Fig. 3C**). Treatment with the LFPRLR SMO also changed the inflammatory T_helper_ (T_h_)1/T_h_17 to a regulatory T_h_2 phenotype (**Figs. 3D and E**), and reduced proliferating Ki67^+^ T_cytotoxic_ cells (**Fig. 3F**) as well as, proliferating Ki67^+^ and potentially pathogenic^34^ CD38^+^CD27^+^CD56^−^CD16^−^ NK cells (**Figs. 3G-H**). LFPRLR knockdown did not induce these changes in the age-and sex-matched healthy donor cells. Thus, in addition to bringing the cellular landscape closer to normal in SLE, there is some initial indication that LFPRLR knockdown may also reduce autoimmune phenotypes that result from other related diseases.

**Fig 3.**
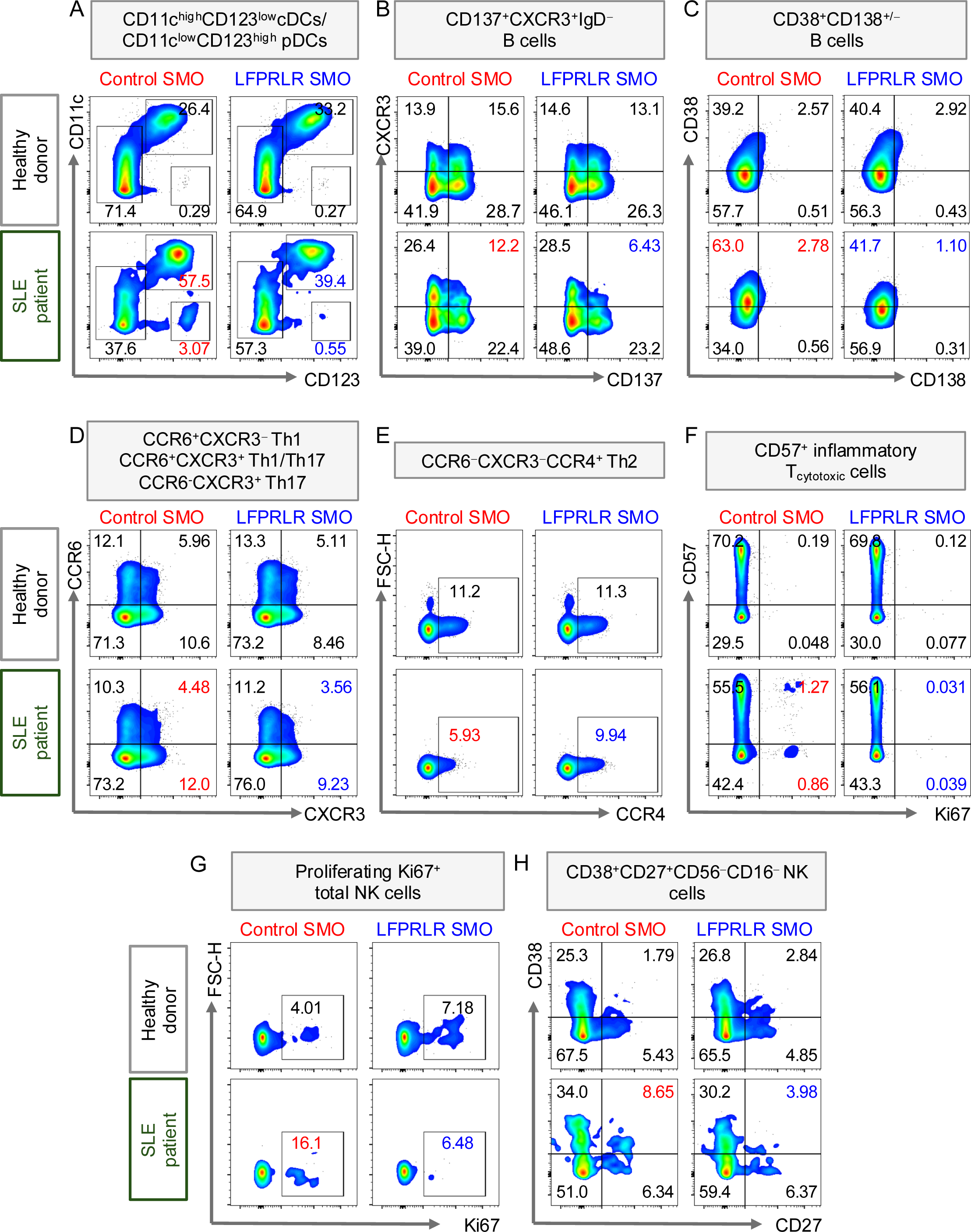
*Ex vivo* knockdown of the LFPRLR in PBMCs of a patient with multiple autoimmune co-morbidities reduced potentially pathogenic dendritic cell (DC), B-cell, T-cell and NK-cell subsets. Flow cytometry depicting percentages of (**A**) CD11c⁺CD123ˡᵒʷ conventional DCs and CD11cˡᵒʷCD123ᵇʳⁱᵍʰᵗ plasmacytoid DCs, (**B**) CD137⁺CXCR3⁺IgD⁻ B cells, (**C**) CD38⁺CD138⁺^/−^ B cells, (**D**) CCR6⁺CXCR3⁺ T_h_1/T_h_17, CCR6⁻CXCR3⁺ T_h_1, and CCR6⁺CXCR3⁻ T_h_17 cells, (**E**) CCR6⁻CXCR3⁻CCR4⁺ T_h_2 cells, (**F**) CD57⁺Ki67⁺ inflammatory cytotoxic T cells, (**G**) Ki67⁺ proliferating total NK cells, and (**H**) CD56⁻CD16⁻CD38⁺CD27⁺ NK cells in PBMC samples from one patient with SLE and age-and sex-matched healthy donor, treated with control SMO or LFPRLR SMO for 72 hours. Pathogenic immune subsets reduced specifically after LFPRLR knockdown in the patient with SLE are depicted in red and blue respectively for control SMO-and LFPRLR SMO-treated patient sample.

Overall, reducing the LF:SF PRLR ratio in cells from patients with SLE concurrently reduces multiple pathogenic immune subsets, bringing them closer to healthy, homeostatic levels. Importantly, this occurs without any impact on cells from healthy donors. No exogenous human PRL was added during the incubations with the SMOs. Thus, the impact of changing PRLR expression on phenotype suggests an important role for autocrine/paracrine prolactin in mediating these effects.

### Reducing the ratio of LF: SF PRLR in SLE patient PBMCs suppresses the expression of type I interferon-stimulated genes

The expression of IFN-I stimulated genes (ISGs) is well known to enhance the generation of autoantibody-secreting plasma cells and exacerbate disease activity in patients with SLE^35^. PRL can induce ISGs^36^. Given that LFPRLR knockdown reduced IFN-I-driven immunophenotypes in SLE (**Figs. 2-3**), we investigated the effects of LFPRLR SMO on the expression of 10 critical ISG transcripts (MX1, MX2, IRF7, OAS1, IFI35, IFI44, IFITM3, STAT1, STAT2, OASL) in PBMCs from three SLE patients with moderate disease activity (SLEDAI 6-10, **Table S1**). We chose these 10 ISGs because (1) their increased expression in blood was found to be associated with increased disease activity, organ damage, spontaneous germinal center (GC) and plasma cell responses, and (2) they have been designated as a biomarker and/or therapeutic target in SLE^37–44^. Hence, we examined the cause-and-effect relationship between the relative levels of PRLR isoforms and the expression of ISGs in SLE patient samples.

*Ex vivo* LFPRLR knockdown for 24 hours significantly reduced the expression of multiple (≥6) ISGs in each of the three patient samples examined (**Fig. 4**). LFPRLR knockdown reduced MX1 expression in all three SLE patient PBMC samples, while the expression of several other genes (MX2, IRF7, OAS1, IFI35, IFI44, IFITM3, STAT1, and STAT2) was reduced after LFPRLR SMO treatment in at least two samples (**Fig. 4**). The simultaneous reduction in multiple pathogenic ISGs after LFPRLR knockdown is concordant with the reduction in the well-known IFN-I-induced pathogenic immunophenotypes observed in **Figs. 2-3**. These results also further corroborate the importance of local PRL, derived from immune cells, in driving pathogenic signaling downstream of LFPRLR in SLE.

**Fig. 4.**
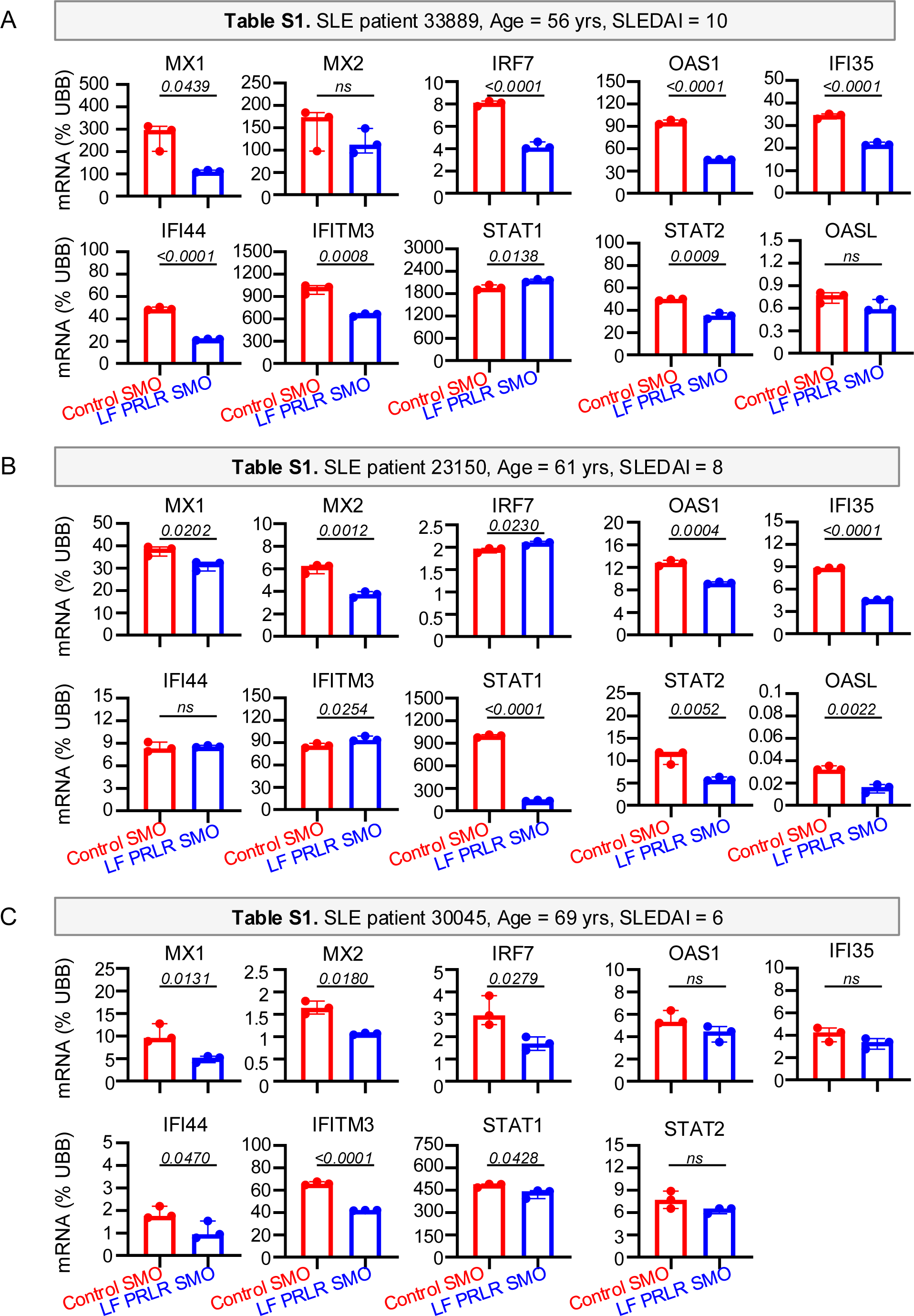
Reducing the ratio of LF: SF PRLR in SLE patient PBMCs suppresses the expression of type I interferon-stimulated genes. (**A-C**) Expression of IFN-I stimulated genes (STAT1, STAT2, MX1, MX2, IRF7, OAS1, OASL, IFI35, IFI44, IFITM3) in PBMCs from an SLE patient cultured *ex vivo* for 24 hours with 500nM of control SMO or LFPRLR SMO. One representative of three independent SLE patient PBMCs is shown. Each dot represents a technical replicate. Transcripts were normalized to UBB and expressed as % of UBB. p-values: Unpaired student’s t-test.

### *Ex vivo* knockdown of the LFPRLR does not impact viability or phenotype of normal humoral immune cells

Mature memory B and plasma cells are increased in pathologies like SLE compared to healthy individuals where few are present in circulation. To determine whether LFPRLR knockdown broadly impacts normal humoral homeostasis mediated by mature human B-cell subsets, we expanded and differentiated the major, mature human B-cell subsets from healthy donor PBMCs *in vitro* (see schema in **Fig. S7A**): IL-4 generates activated non-switched IgD^+^CD27^+^ memory B cells, while IL-21 drives the production of switched IgD^−^CD27^+^ memory B cells; IgD^−^CD27^−^ extrafollicular, atypical B cells; and CD138^+^CD38^bright^ plasma cells (**Figs. S7B-C**).

Knockdown of the LFPRLR *in vitro* for 48 hours did not affect the viability or phenotype of activated, memory, or plasma B-cell subsets expanded from healthy donor PBMCs, even in the presence of added exogenous PRL that mimics elevated circulating PRL in SLE (**Fig. S8**). Thus, LFPRLR knockdown does not: (1) broadly lead to immunosuppression, as is seen in the case of steroid and other immunomodulatory treatments for SLE, or (2) alter humoral homeostasis, as is seen in the case of B/plasma cell-depleting therapies.

### *In vivo* LFPRLR knockdown in SLE-prone mice reduces IFN-I signaling and response in immune cells

To address the potential long-term benefits of LFPRLR knockdown, we administered the LFPRLR SMO for 8 weeks at a dose of 100 pmole/hour/mouse to 6-week-old *MRL-lpr*^45^ primary SLE-prone mice (**Fig. 5A**). By preventing the death of autoreactive immune cells, the *Fas^lpr^* mutation in these mice results in morbidity from systemic autoimmunity in 17-22 weeks^45^. Although in a study of B-cell transformation^14^, we have recently reported an 8-week LFPRLR knockdown to reduce the number of splenic B cells, detailed characterization of pathogenic immunophenotypes of PRLR^+^ and PRLR^−^ B-and non-B cells after such knockdown was not performed. We therefore conducted scRNA-seq to compare transcriptional changes in both PRLR^+^ and PRLR^−^ immune cells in healthy mice (n=3) and in SLE-prone mice treated with control SMO-(n=4) or LFPRLR SMO (n=4) (**Fig. S9** for protocol).

**Fig 5.**
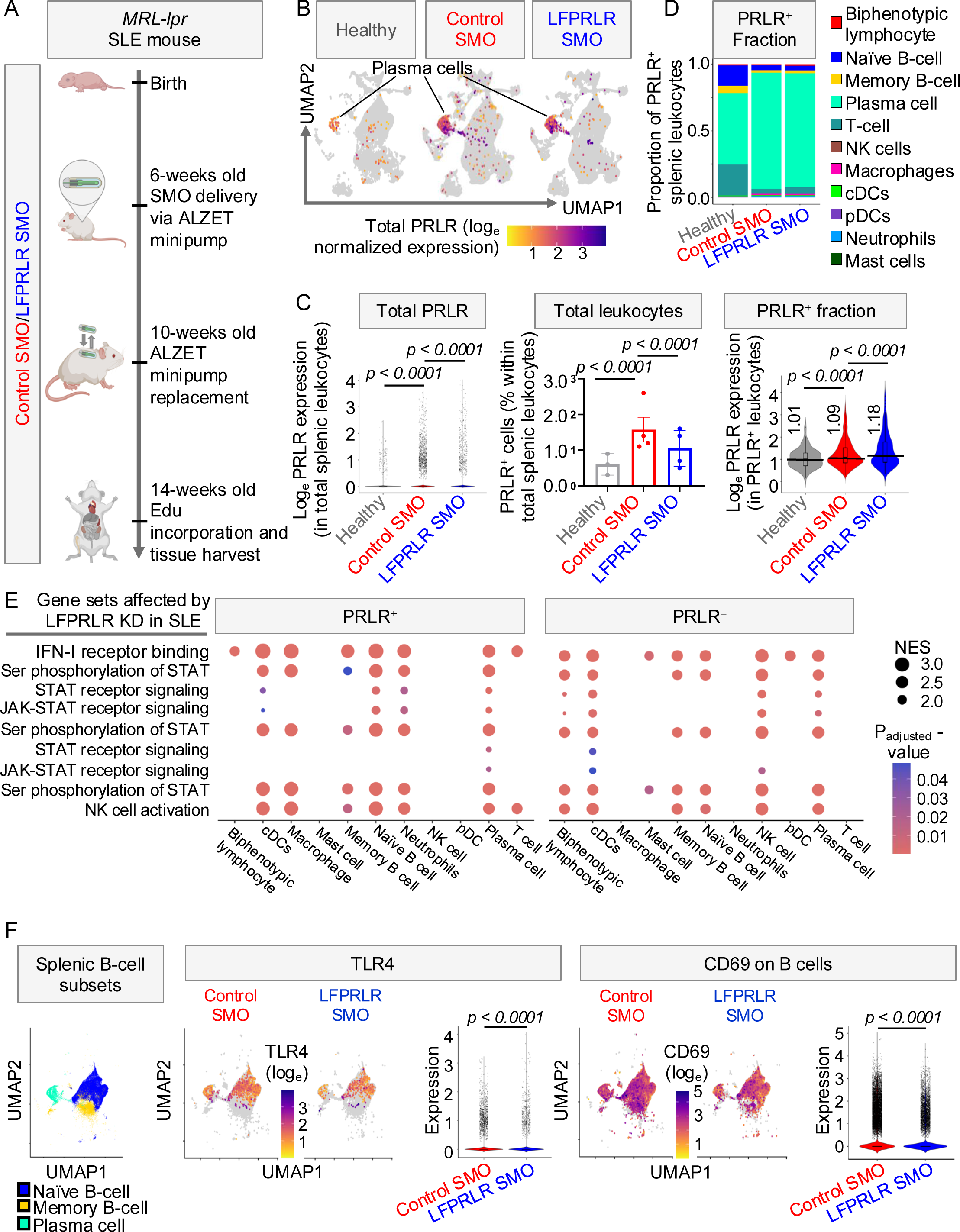
*In vivo* LFPRLR knockdown in SLE-prone mice reduces IFN-I signaling and response in immune cells. (**A**) Experimental approach to measure the *in vivo*, long-term effects of LFPRLR knockdown on immunopathology in primary *MRL-lpr* SLE-prone mice. (**B**) Expression of total PRLR (predominantly LF, not shown) in spleens of healthy mice (n=3), and control SMO-(n=4) and LFPRLR SMO-(n=4) treated *MRL-lpr* SLE-prone mice. Publicly available healthy C57BL/6 splenic scRNA-seq datasets were downloaded from 10X Genomics. (**C**) Overall distribution of PRLR expression in leukocytes (left), proportion of PRLR^+^ leukocytes (middle, mean ± SEM), and expression of PRLR transcripts within PRLR^+^ leukocytes (right) in spleens of healthy mice (n=3), and control SMO-(n=4) and LFPRLR SMO-(n = 4) treated *MRL-lpr* SLE-prone mice. P-values for left, middle, and right panels in (C) have been calculated by t-test for bimodal distribution, chi-squared, and Mann-Whitney U test respectively. (**D**) Distribution of immune cell types in the PRLR^+^ splenic leukocyte fraction of healthy mice (n=3), control SMO-treated (n=4), and LFPRLR SMO-treated (n=4) *MRL-lpr* mice by scRNA-seq. (**E**) Gene ontology enrichment analysis by scRNA-seq showing the topmost differentially regulated pathways in each splenic leukocyte subset of *MRL-lpr* SLE-prone mice treated with control SMO (n=4) or LFPRLR SMO (n = 4) *in vivo*. (**F**) Comparison of transcript expression of TLR4 and B-cell activation marker CD69 in splenic B-cell subsets (naïve, memory, and plasma) of *MRL-lpr* SLE-prone mice treated with control SMO (n = 4) or LFPRLR SMO (n=4) *in vivo*. p-value: Seurat’s implementation of the likelihood ratio test for bimodal distribution.

Although *MRL-Mpj* is the usual control for the *MRL-lpr* model, these mice will eventually develop SLE, albeit at a slower rate than their *MRL-lpr* counterparts. To model the immune system in an entirely normal mouse/healthy mouse we used *C57BL/6J*. The proportion of PRLR^+^ cells and expression of PRLR in PRLR^+^ cells were both aberrantly increased in SLE compared to healthy mice. LFPRLR knockdown brought the proportion of PRLR^+^ splenic leukocytes in SLE-prone mice closer to those seen in normal mice (**Figs. 5B-C**). LF was the predominant PRLR isoform in splenic immune cells. However, after the 8-week treatment period, the LF:SF ratio was unaffected by LFPRLR knockdown (**Fig. S10**), suggesting that treatment with LFPRLR SMO for 8 weeks may have already eradicated splenocytes with reduced LFPRLR by a Fas-independent mechanism.

Within the PRLR^+^ splenic fraction in SLE-prone mice, PRLR expression was highest in the splenic plasma cells (**Figs. 5B, 5D, S10**), although most plasma cells were PRLR^−^. PRLR^+^ splenic plasma cell fraction was increased in SLE compared to healthy mice (**Fig. 5D**). In addition, macrophages and neutrophils in SLE-prone mice aberrantly expressed the PRLR (**Figs. 5D, S11**). Thus, a greater array of immune cells is likely sensitive to PRL in SLE-prone mice compared to their healthy counterparts.

There were no major changes in the relative distribution of PRLR^+^ immune subsets after LFPRLR knockdown (**Figs. 5D** and profiles in **S10**). However, LFPRLR knockdown significantly impacted pathways associated with JAK-STAT and IFN-I signaling, and NK-cell activation in PRLR^+^ and/or PRLR^−^ immune cell subsets (**Fig. 5E**), indicating both direct and multiple indirect effects of the knockdown. Consistent with this, transcription of multiple STATs (1,2,3,5a,5b) was reduced after LFPRLR knockdown, and significant reductions were seen in the numbers of STAT^+^ plasma cells and macrophages (**Figs. S12-13**).

Transcripts of toll-like receptor 4 (TLR4) (**Fig. 5F**), TLR4 downstream signaling molecules (**Fig. S14**), and CD69 (**Fig. 5F**), all of which are indicative of TLR4-induced IFN-I signaling and B-cell activation^46^, were reduced in B-cell subsets of SLE-prone mice after LFPRLR knockdown. Thus, LFPRLR knockdown suppresses pathogenic signaling that would otherwise lead to the activation of autoreactive B-cell subsets in secondary lymphoid tissues such as the spleen.

### *In vivo* LFPRLR knockdown in SLE-prone mice reduces the production and proliferation of plasma cells and brings the transcriptional signature of GC and transitional B cells closer to that seen in healthy mice

Even though there were no major changes in PRLR^+^ immune subsets after LFPRLR SMO treatment, as mentioned above, knockdown significantly reduced numbers of both splenic PRLR^+^ and PRLR^−^ plasma cells (**Fig. 6A**), suggesting that LFPRLR SMO can eradicate plasma cells both directly and indirectly. Of interest therefore was whether the reduction in plasma cells was due to increased apoptosis, reduced proliferation, and/or reduced generation. There was no effect of LFPRLR knockdown on expression of genes that define an apoptotic signature^47^ (**Fig. S15**). However, LFPRLR knockdown reduced Ki67^+^ proliferating PRLR^+^ and PRLR^−^ plasma cells (**Fig. 6B**). Based on LFPRLR’s role in driving TLR4/IFN-I-induced signaling and B-cell activation in SLE (**Figs. 5E-F, S14**), we hypothesized that LFPRLR knockdown would block the generation of autoantibody-secreting plasma cells. Plasma cells develop from both splenic follicular (FO) B cells, and those in the extrafollicular marginal zone (MZ). While FO B cells undergo affinity maturation in splenic germinal centers (GCs) and give rise to class-switched (IgM/IgD⁻) memory B cells and plasma cells, MZ B cells produce non-class switched memory B and plasma cells^48^. Knockdown of LFPRLR in SLE-prone mice led to a decreased frequency of FO B cells and a concomitant increase in MZ B cells (**Fig. 6C**, gating strategy in **Fig. S16**).

**Fig. 6.**
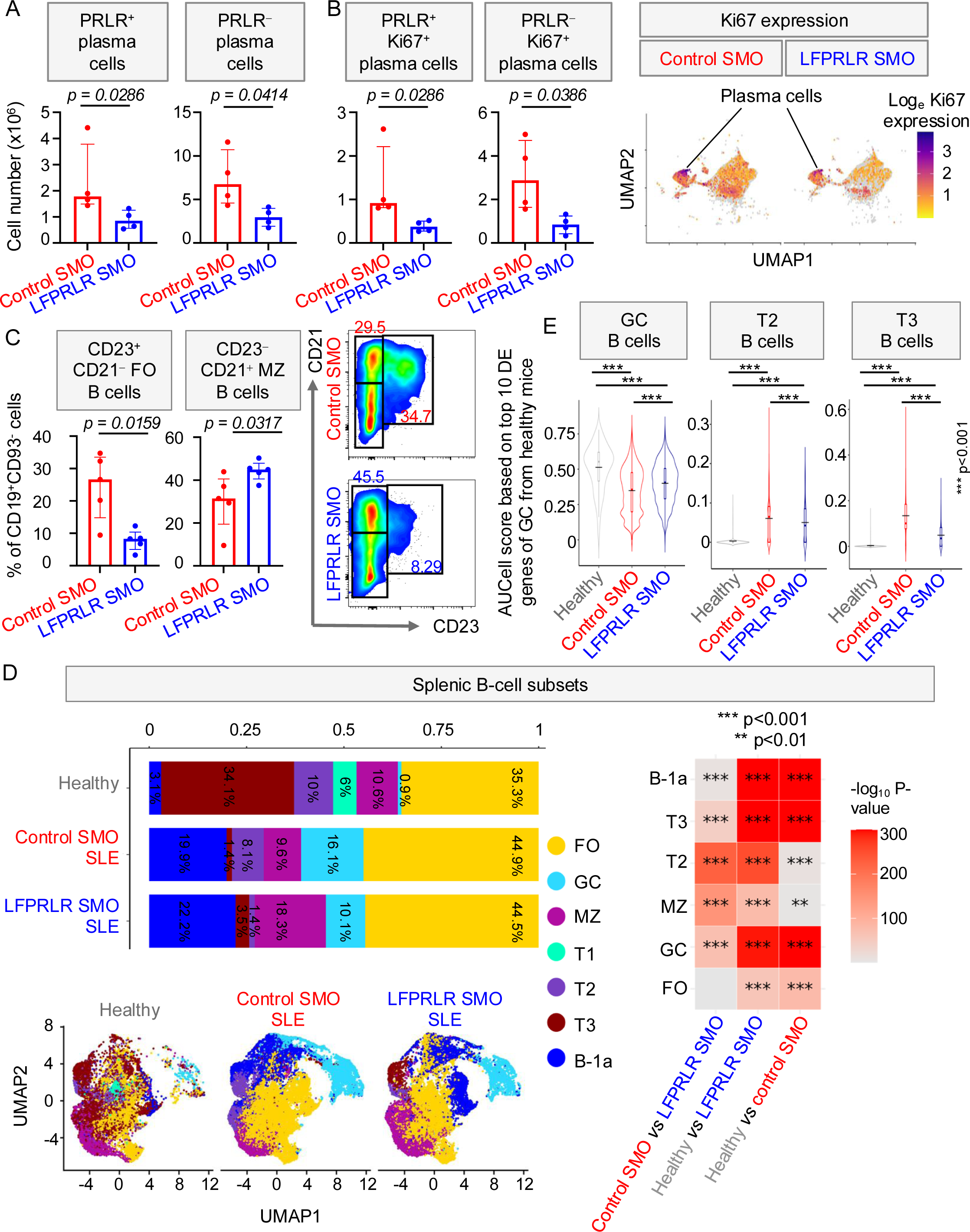
*In vivo* LFPRLR knockdown in SLE-prone mice reduces the proliferation and production of plasma cells and brings the transcriptional signature of GC and transitional B cells closer to that seen in healthy mice. (**A-C**) Effects of *in vivo* knockdown of the LFPRLR in *MRL-lpr* mice on (**A**) numbers of PRLR⁺ and PRLR⁻ splenic plasma cells by scRNA-seq, (**B**) numbers of Ki67⁺ splenic plasma cells within PRLR⁺ and PRLR⁻ populations by scRNA-seq, (**C**) CD23⁺CD21⁻ follicular (FO) and CD23⁻CD21⁺ marginal zone (MZ) B cells within the CD19⁺CD93⁻ splenic B cell fraction by flow cytometry. Data are shown for scRNA-seq and flow cytometry in control SMO-and LFPRLR SMO-treated arms from 4 and 5 *MRL-lpr* SLE-prone mice per group, respectively. Median ± interquartile range, p-values: Mann-Whitney U test. (**D**) Distribution of splenic B-cell subsets in healthy mice (n=3), and control SMO-(n=4) and LFPRLR SMO-(n=4) treated *MRL-lpr* SLE-prone mice. P-values: Chi-square test. (**E**) AUCell score of GC, T2, T3 splenic B cell subsets in healthy mice (n=3), and control SMO-(n=4) and LFPRLR SMO-(n=4) treated *MRL-lpr* SLE-prone mice. AUCell score was calculated based on the top 10 differentially expressed (DE) genes for each subset derived from healthy mice. Median ± interquartile range (black line depicts mean of each group), p-value: Mann-Whitney U test.

Using scRNA-seq, we further characterized the effect of LFPRLR knockdown on splenic B cells by detailed transcriptional profiling of each major B-cell subset [FO, MZ, GC, B-1a, and transitional (T1-T3)]. In accordance with increased plasma cell production in SLE, transitional cells T1-T3 were reduced in SLE-prone mice compared to healthy ones. LFPRLR knockdown in SLE-prone mice increased the proportion of T3 but, at least with the dose and duration of treatment examined, could not restore to levels seen in healthy counterparts. Consistent with **Fig. 6C**, the proportion of B cells with MZ transcriptional signature was increased after LFPRLR knockdown in SLE-prone mice, whereas those with GC-like transcriptomic profile were reduced (**Fig. 6D**). Reduced GC-like cells after LFPRLR knockdown were substantiated by reductions in IgM/IgD⁻ class-switched memory B cells and plasma cells, and in numbers of CD11c^−^CD11b^−^ B cells, shown recently^49^ to exhibit an activated plasmablast signature (**Figs. S17A-B**). The age-associated, potentially extrafollicular CD11c^+^CD11b^−^ B-cell subset^49^ was also reduced after LFPRLR knockdown in SLE-prone mice (**Fig. S17B,** gating strategy in **Fig. S18**).

To determine the extent to which LFPRLR knockdown restores B-cell homeostasis in SLE, we then examined the similarity between the gene expression signatures of the different B-cell subsets in healthy mice, control SMO-and LFPRLR SMO-treated SLE-prone mice using the AUCell method^50^. Notably, LFPRLR knockdown in SLE-prone mice partially brought the pattern of expression of genes in GC, T2, T3, and B-1a B cells closer to that seen in healthy mice (**Figs. 6E**, **S19B**; heatmaps of genes used to calculate the AUC score for each subset are provided in **Fig. S19A**). Among the many changes in gene expression across B-cell subsets shown on the heatmap, the GC fraction was particularly interesting. Many of the highly expressed genes in GC cells in healthy animals (**Fig. S19A**, healthy cells, second column) are reduced in SLE (**Fig. S19A**, control SMO SLE, second panel) and partially restored by LFPRLR knockdown (**Fig. S19A**, LFPRLR SMO SLE, second panel). Among these, it is interesting to note Tent5c, which is a non-canonical cytoplasmic poly(A) polymerase highly expressed by activated B cells that suppresses their proliferation^51^, and syndecan 1 (sdc1, CD138), which is a cell surface heparan sulfate proteoglycan that binds to many soluble mediators of inflammatory disease^52^. Overall, our results suggest that increased LF:SF PRLR expression in SLE disturbs the homeostasis of B-cell subsets and drives the formation of B cells with GC-like transcriptional signatures.

### *In vivo* LFPRLR knockdown in SLE-prone mice reduces expression of splenic B cell immunoglobulins with features of autoreactivity

Recent studies from us and others have shown the importance of PRL signaling through the LFPRLR in driving the production of anti-double-stranded DNA autoantibodies^12,15^. However, the extent to which changes in PRLR isoforms (increased LF:SF) in SLE more broadly impacts B/plasma cells that exhibit autoreactive signatures had not been investigated.

In humans, antibodies more likely to exhibit autoreactivity possess complementarity determining region 3 (CDR3) lengths exceeding 15-20 amino acids (aa)^53^, are class-switched for greater affinity^54^, and have a characteristic amino acid composition enriched in charged residues^55^.

We now describe for the first time that, as in humans, SLE-prone mice have antibodies with CDR3≥15aa that possess an amino acid composition similar to that seen in human autoantibodies: clustered heatmaps of aa usage show different frequencies between CDR3>15aa versus CDR3 <15aa in total, memory, naïve B, and plasma cells (**Fig. 7A-D**). Importantly, in SLE-prone mice, treatment with the LFPRLR SMO reduced the CDR3aa length in total, naïve, and memory B cells and appeared to do so in plasma cells, although this did not reach statistical significance (**Fig. 7E**). Focusing on plasma cells from the SLE-prone mice, one can see that cells producing antibodies with CDR3>15aa are more frequently class switched than cells producing antibodies with CDR3<15aa (only 3.5% are not class switched in CDR3>15 *versus* 18.1% in CDR3<15) (**Fig. 7F**). Importantly, the proportion of non-class switched antibodies is increased in response to LFPRLR SMO, regardless of CDR3 length. Said another way, the proportion of plasma cells producing potentially autoreactive antibodies is reduced with LFPRLR SMO. Therefore, unlike agents such as belimumab, which reduce activated, autoreactive B cells without affecting pathogenic CDR3 features^55^, LFPRLR SMO can reduce both.

**Fig. 7.**
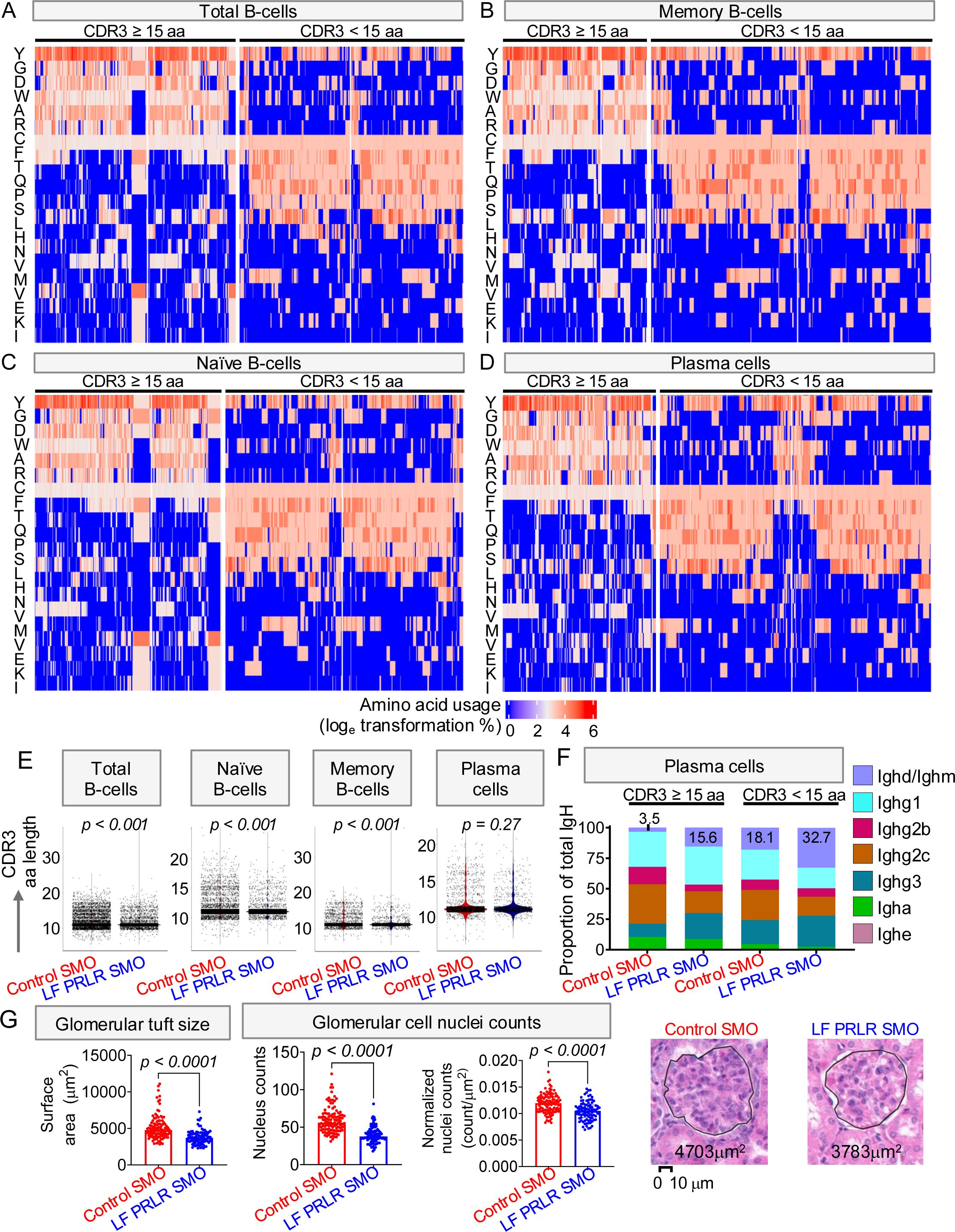
*In vivo* LFPRLR knockdown in SLE-prone mice reduces splenic B cell immunoglobulins with features of autoreactivity and prevents the development of glomerular pathology. (**A-D**) Heatmaps comparing aa sequences of IgH CDR3 ≥ 15aa and IgH CDR3 < 15aa in pan B cells and within specific B-cell subsets in *MRL-lpr* SLE-prone mice (n=4 control SMO, n=4 LFPRLR SMO) by scRNA-seq; and (**E-F**) scRNA-seq VDJ analysis in SLE-prone *MRL-lpr* mice treated with control SMO (n=4) or LFPRLR SMO (n=4) for 8 weeks *in vivo*. (**E**) distribution of immunoglobulin heavy chain (IgH) CDR3 length in amino acids (aa) in total B cells and all B-cell subsets. (**F**) IgH isotype within long and short CDR3-containing plasma cell subsets. P-values: Mann-Whitney U test; (**G**) Representative H&E images and quantitation of glomerular tuft size and mesangial cell nuclei counts, both raw and normalized, in control SMO-and LF PRLR SMO-treated *MRL-lpr* mice. Bar plots: median ± interquartile range, p-values: Mann-Whitney U test.

### *In vivo* knockdown of LFPRLR averts development of glomerular pathology in SLE-prone mice

The kidneys of SLE-prone mice were examined at 14 weeks of age after 8-week LFPRLR knockdown and compared to control SMO-treated animals. In lupus, glomerular tufts, which are the blood filtering units in the kidney, aberrantly increase in size, cellularity, and cellular density (cells per unit area), due in part to immune cell infiltration and in part to mesangial cell proliferation^56–58^. Analysis of glomerular cellularity is appropriate to this stage of disease, which is prior to significant basal laminar thickening or crescent formation. Knockdown of LFPRLR averted the development of hypercellularity in the glomeruli (**Fig. 7G**), demonstrating that this approach effectively prevents glomerular damage. At 14 weeks of age in the SLE-prone mice, glomerular function is just beginning to be disturbed. In control SMO-treated animals, 3 mice had developed significant proteinuria (determined by outlier analysis in **Fig. S20**). There were no mice with outlier values in the LFPRLR SMO treatment group. In semi-quantitative analysis, LFPRLR SMO treatment of the SLE mice reduced the mean value for calciuria but this reduction did not reach statistical significance compared to the control group (**Fig. S20**).

## Discussion

While clinical evidence has associated elevated circulating PRL with exacerbation of SLE, little was previously known about how PRL affects the course of this disease. Using primary human samples and SLE-prone mice, we are the first to show that there is aberrant PRLR splicing in multiple immune subsets in SLE. Further, this aberrant splicing produces an increased ratio of LF:SF PRLR. Using a splice modulating oligomer that reduces expression of the LFPRLR and/or increases expression of the SFPRLR, we demonstrate through at least partial normalization of multiple pathogenic immune subsets that the change in LF:SF ratio is causal in disease. The immune subsets examined included both self-reactive B cells and other immune cells like macrophages, NK cells, and cDCs that can fuel B-cell activation. Interestingly, both PRLR^+^ and PRLR^−^ cells were responsive, indicating that in addition to direct effects, knockdown of the LFPRLR caused changes that affected other cells in the environment, presumably by the secretion of local paracrine cytokines. Such mechanisms would be subjects for future investigations.

In the course of this work, we also documented the breadth of immunologic sources of autocrine/paracrine PRL. In doing so, we explain why some patients might not substantially benefit from simply reducing pituitary PRL output with dopamine agonists since dopamine agonists only inhibit the secretion of pituitary and not extrapituitary PRL^4,5^. The importance of immune cell autocrine/paracrine PRL to the progression of disease is demonstrated by the beneficial effects of LFPRLR knockdown in the *ex vivo* experiments where no PRL was added to the medium.

The LFPRLR SMO alters splicing by preventing the inclusion of exon 10 into the mature mRNA. In many cells, this just reduces expression of the LFPRLR^14^. However, in some cells, where the difference is presumably governed by different splicing factor expression, reducing inclusion of exon 10, promotes expression of the SFPRLRs^15^ (almost exclusively SF1b in human immune cells shown herein). Previous studies have demonstrated that increased expression of LFPRLR results in proliferation and/or survival, whereas increased expression of SFPRLRs inhibits proliferation and promotes differentiation and/or apoptosis, depending on the relative ratios^4^. Specific knockdown of the LFPRLR, which may be accompanied by upregulation of the SFPRLRs, allows for continued signaling through the SFPRLRs^59^ and includes signaling by SF1b that results in down regulation of expression of the LFPRLR^60^. Thus, not only is the signal to proliferation/survival reduced, but the signal to differentiation and/or apoptosis is continued/increased^59,60^. This is a significant advantage that would not occur with PRLR antagonists or anti-PRLR antibodies^61^ or anti-PRL antibodies, each of which would target all splice forms of the PRLR that have identical extracellular domains. Thus, more traditional receptor targeting strategies would remove all beneficial signaling through SFs at the same time as the pathogenic signaling through excess LF.

Importantly, we also show that isoform-specific knockdown of LFPRLR had no discernible negative impact on normal immune cells. This is, to our knowledge, the first report of an effective strategy for SLE treatment that only affects pathogenic cells. Further, based on studies in normal mice, which lasted for 18 months and showed no signs of toxicity, the LFPRLR SMO is very well tolerated^14^.

Finally, to determine whether LFPRLR knockdown and associated changes in pathogenic immune cell subsets could substantially alter a long-term consequence of SLE, we examined the development of glomerular pathology and found that knockdown of the LFPRLR averted development of characteristic hypercellular, histopathological changes. Thus, it appears that the knockdown of the LFPRLR not only largely normalizes pathogenic immune subsets but also inhibits the development of a major cause of morbidity and mortality in this disease. Thus, LFPRLR SMO has significant potential as an effective therapeutic agent in SLE that may be more easily tolerated than what is currently clinically available^1–3^. Considering agents currently in development and undergoing clinical trials for SLE, autologous CD19-specific chimeric antigen receptor (CAR) T cells have shown promise in patients with active SLE^62^. However, unlike LFPRLR SMO, CAR T cells can only be administered in inpatient settings at specialized centers, are costly, and were recently shown to have minimal impact on autoantibody profiles of patients^63^. CAR T cell therapy may result in cytokine release syndrome and mild immune effector cell associated neurotoxicity syndrome^64,65^. LFPRLR SMO shows no signs of either syndrome^14^, likely because SFPRLRs are expressed in the brain, and PRL acting through SFPRLR results in neuroprotection^66^.

Of additional interest is our recent determination of the role of LFPRLR in driving the evolution of B-cell lymphomas^14^ since the incidence of lymphoma is 3-8-fold higher in patients with SLE and related autoimmune diseases^67–69^. Because clinical investigations in patients with SLE suggest that autoreactive and malignant processes proceed in parallel and are likely to occur in different B cells^67,68^, we predict that the LFPRLR SMO could be a promising approach for treating autoimmunity and simultaneously lowering the likelihood of development of lymphoma in patients with SLE.

SLE is nine times more frequent in adult females than males^7,70,71^, and hence our examination of LFPRLR knockdown sequelae was conducted in cells from adult female patients and female SLE-prone mice. As mentioned previously ^reviewed^ ^in^ ^6^, estrogen stimulates the production of pituitary PRL, adult females have higher circulating levels of PRL than males or prepubertal children, and postmenopausal women treated with hormone therapy have been found to have an increased risk of developing SLE^72^. However, SLE does occur in male patients, in postmenopausal women and children of both sexes^73,64^ and, when it occurs, tends to be a more severe disease. The regulation of PRL production in extrapituitary sites is more complex than in the pituitary and likely shows some differences among cell types^74^. However, as an example, stress elevates PRL, and epinephrine has been shown to increase monocyte/macrophage production of autocrine/paracrine PRL^75^. It may be this local elevation of immune cell PRL that makes the disease more severe.

A limitation of our study was our inability to procure treatment-naïve SLE patient samples. Prior treatment likely added to the degree of variability observed in response to LFPRLR knockdown. Nevertheless, our findings clearly show that LFPRLR overexpression, without concomitant increases in SFPRLR expression, drives SLE and that specific LFPRLR knockdown represents a viable strategy to normalize immune subsets and avert glomerular pathology. With the recent FDA approval of other splice-modulating oligonucleotide therapies^76^, development of LFPRLR SMO as a therapeutic for SLE seems justified in light of its multiple benefits over current therapeutic regimes. However, regardless of whether LFPRLR SMO makes it through to clinical use, its experimental use has uncovered several mechanisms through which PRL-PRLR interactions contribute to the development of SLE and opened up many interesting avenues for further investigation.

## Materials and Methods

### SMO

Human and mouse LFPRLR SMOs (sequences for these and control SMO provided in^15^ and in **Table S2a**) are 25mer morpholino DNA oligos that block the inclusion of exon 10, essentially specific to LF/IF PRLRs during pre-mRNA splicing^14,15,77^. The SMOs are linked to an octaguanidine dendrimer that ensures effective whole-body uptake and are administered subcutaneously (see below). They were custom synthesized by GeneTools LLC (Philomath, OR, USA). Unlike small interfering RNAs (siRNAs), they do not require nanocarrier delivery. We have previously shown that healthy mice treated with the LFPRLR SMO for 18 months showed no detectable toxicity, as assessed by behavior, weight, liver enzymes, blood cell counts, and tissue histology^14,77^.

### Healthy donor and SLE patient samples

Deidentified peripheral blood samples from SLE patients (**Table S1**) were procured from Sanguine Biosciences (Woburn, MA, USA). Peripheral blood samples from age-and sex-matched healthy donors were procured from the Michael Amini Transfusion Medicine Center of City of Hope (Duarte, CA, USA). Peripheral blood mononuclear cells (PBMCs) were isolated by Ficoll-Hypaque density gradient separation (Cytiva, Marlborough, MA, USA) and cryopreserved in liquid nitrogen using 10% dimethyl sulfoxide (DMSO) in fetal bovine serum (FBS). This study does not involve direct contact with human subjects and is classified as non-human subjects research under City of Hope Institutional Review Board (IRB) 19373.

### Comparison of LFPRLR knockdown-induced immunophenotypes in healthy human and SLE patient PBMCs

Cryopreserved human PBMCs were thawed and cultured in RPMI-1640 medium (Invitrogen/Life Technologies, CA, USA) supplemented with 10% FBS and 1% penicillin and streptomycin (termed ‘complete RPMI’). Cells were plated in a 96-well U-bottom plates at a concentration of 2×10^6^ cells/mL and treated with either control SMO or human LFPRLR SMO for 72 hours at a concentration of 500nM. This concentration was selected based on preliminary dose-response experiments performed to define the dynamic range for *ex vivo* use in human PBMCs ^15^. Following culture, cells were harvested and prepared for flow cytometry analysis.

Changes in NK and monocyte phenotype after LF PRLR knockdown were determined as ratios of LFPRLR SMO-induced immunophenotype versus control SMO-induced immunophenotype for each sample. For both NK and monocyte fractions, we calculated the relative proportions of the less mature (CD56^bright^ for NK and CD14⁺CD16⁻ for monocytes) and more mature (CD56^dim^ for NK and CD14⁺CD16^+^ for monocytes) subsets. To compute the immunophenotypic effects, we used the formula: (relative proportion of less mature to more mature subset in LFPRLR SMO) ÷ (relative proportion of less mature to more mature subset in control SMO).

### High-dimensional flow cytometry

Cell viability was assessed using LIVE/DEAD^TM^ (Thermo Fisher Scientific, Waltham, MA, USA) for 20 minutes at 4°C. Surface staining was performed by incubating cells for 30 minutes at room temperature. For intracellular staining, cells were fixed and permeabilized using eBioscience™ Foxp3/Transcription Factor Staining Buffer Set (Thermo Fisher Scientific, Waltham, MA, USA) according to the manufacturer’s instructions, followed by intracellular antibody staining for 30 minutes at room temperature. Antibody dilutions were prepared according to the manufacturer’s recommendations and optimized in preliminary experiments. All antibodies (**Table S3-S4**) were obtained from BioLegend, BD Biosciences, or Thermo Fisher Scientific. Data acquisition was performed on Cytek Aurora (Cytek Biosciences, Fremont, CA, USA) or BD FACSymphony (BD Biosciences, San Jose, CA, USA) cytometers and analyzed using Flowjo software (version 10.7.1; BD Biosciences, OR USA). Gating strategies for human PBMC immune cell populations are detailed in **Fig. S4**. The gating strategy for murine splenic immune cells was described in an earlier publication ^15^.

### Quantitative real-time PCR (qPCR)

RNA was extracted using the RNAaqueous™ Micro kit (Micro Scale RNA Isolation Kit; Thermo Fisher Scientific, Waltham, MA, USA) according to the manufacturer’s instructions. cDNA was synthesized using SuperScript™ IV Reverse Transcriptase (Invitrogen, Carlsbad, CA, USA). qPCR was performed using Power SYBR Green PCR Master Mix (Thermo Fisher Scientific, Waltham, MA, USA) on a QuantStudio 7 Flex real-time PCR system (Applied Biosystem, Foster City, CA, USA). qPCR primer sequences are listed in **Table S2b**.

### Animal models and *in vivo* administration of SMOs

Animal studies were conducted in compliance with the Institutional Animal Care and Use Committee (IACUC #19032, #23150) at the City of Hope. 6-week-old female SLE-prone *MRL-lpr* mice received either control SMO or mouse LFPRLR SMO (sequences of SMO are in Table S2a^15^) via subcutaneously implanted Alzet minipumps (Durect Corporation, CA, USA) for 8 weeks, as described previously^15^. Alzet pumps delivering 100 pmoles/h/mouse were replaced at the 4^th^ week of treatment. Two hours before euthanasia and after 8 weeks of treatment, each animal received an intraperitoneal injection of 2.8 mg of the nucleoside analog, 5-ethynyl-2′-deoxyuridine (EdU, Invitrogen, Carlsbad, CA, USA) in PBS.

### Mouse tissue processing

As described in our previous study^15^, splenic leukocytes were isolated from *MRL-lpr* mice and stored in liquid nitrogen as frozen single cell suspensions in 90% FBS and 10% DMSO until analysis. Kidneys were fixed for at least 24 hours in 4% formaldehyde (diluted from buffered 37% formaldehyde, Thermo Fisher Scientific, Waltham, MA, USA). The fixed kidneys were transferred to 70% alcohol. Further dehydration, clearing, and paraffin infiltration were performed on a Tissue-Tek VIP 6 AI Vacuum Infiltration Tissue Processor (Sakura Finetek, CA, USA). The tissues were embedded in paraffin wax using a Tissue-Tek TEC Tissue Embedding Station (Sakura Finetek, CA, USA) prior to sectioning and hematoxylin and eosin (H&E) staining.

### Hematoxylin and eosin (H&E) staining

Sectioned kidney slides were stained on a Tissue-Tek Prism Plus Automated H&E Stainer (SAKURA) according to standard laboratory procedures. Whole slide microscope images were acquired with a Ventana iScan HT Scanner (Roche Diagnostics, IN, USA) at City of Hope Research Pathology Core.

### Analysis of kidney glomerular pathology

H&E-stained kidney ndpi images were imported into the NDP viewer (Hamamatsu Photonics, Hamamatsu City, Japan). Because lupus pathology can be regional, each coronal kidney section was partitioned into three regions (top, middle, and bottom). After determining that the total number of glomeruli did not differ between groups at this relatively early stage of pathology, Three representative glomeruli were selected in each region of both kidneys, for a total of 18 glomeruli per animal. After controlling for potential errors due to the glomerular section plane, the area of each glomerular tuft in the section was measured using the NDP viewer’s outlining tool, and nuclei within each glomerulus were counted. Normalization per unit area was performed by dividing the total number of nuclei per glomerulus by the area of the respective glomerulus.

### Urinary parameters

Urine was collected by placing mice in small chambers over sterile 96-well plates and waiting for natural micturition. Samples were transferred to sterile tubes and frozen until batch analysis. Total urinary protein was measured by Bradford assay using bovine serum albumin as the standard protein. Calciuria was assessed in duplicate with colorimetric multi-analyte urine test strips (Complete Natural Products, Wood Cross, UT, USA).

### Calculation of absolute numbers of splenic leukocytes in SLE-prone mice

Total leukocytes in splenic single cell suspensions were counted using a hemocytometer before subjecting these cells to downstream analysis including single-cell (sc) RNA-seq. The proportions of specific PRLR^+^ and PRLR^−^ immune subsets were determined by scRNA-seq and, in conjunction with total splenic leukocyte counts, were used to calculate the absolute numbers of PRLR^+^ and PRLR^−^cells within each immune subset.

### Human mature B cell production protocol

After thawing PBMCs at 37°C, they were centrifuged at 350 × g for 10 min, washed twice with MACS buffer (DPBS + 0.5 mM EDTA), and counted using a hemocytometer. A 100 μL aliquot was set aside for flow cytometry. Remaining cells were labeled with biotin-antibody cocktail and anti-biotin MicroBeads (Miltenyi Biotec, Bergisch Gladbach, Germany) per manufacturer’s instructions. Pan B cells were isolated using LS Columns and counted; 50–100 μL was reserved for phenotyping by flow cytometry.

The remaining cells were cultured in StemMACS HSC Expansion Medium (no cytokines) at 4×10⁶ cells/mL. Parallel cultures were supplemented with CD40 ligand (CD40L) together with IL-4 or IL-21. Media were refreshed on days 2 and 4 of treatment (with CD40L+IL-4 or CD40L+IL-21). M2-10B4 feeder layer cells (American Type Culture Collection, VA, USA) were plated and adhered to a 0.2% gelatin-coated plate on day 5. On day 6, the M2-10B4 feeder layer medium was removed and replaced with complete RPMI medium with 10μg/mL mitomycin-C (Sigma-Aldrich, MO, USA) for 3 hours. The feeder layer cells were then washed with DPBS and cultured with either CD40L+IL-4 or CD40L+IL-21 media. The B cells were restimulated and then plated in Transwell™ inserts that were placed on top of the mitomycin-C-treated feeder cells.

Restimulation with CD40L+IL-4 or CD40L+IL-21 occurred on days 9, 14, 16, 20, and 21. Viability, surface immunophenotype by flow cytometry, and cell counts were measured on days 0, 13 and/or 14, and 21 and/or 23. On day 14, IL-4 and IL-21 cultures were analyzed by flow cytometry (antibodies in **Table S5**) and 3-(4,5-dimethylthiazol-2-yl)-5-(3-carboxymethoxyphenyl)-2-(4-sulfophenyl)-2H-tetrazolium salt (MTS) assay. To induce further plasma cell production, the remaining IL-4-treated cells were switched to IL-21 + CD40L on day 13 or 14, cultured for 7-9 more days on M2-10B4 feeder cells, and finally analyzed by flow cytometry and MTS assay.

### MTS cell fitness assay

Twenty-five thousand cells per well were seeded in 96-well plates and incubated in Complete RPMI containing control or LFPRLR SMO (0, 5, 25, 125, 250, 500, 750, 1000, 1500, 2000 nM) and 200ng/mL exogenous human recombinant PRL (PeproTech, NJ, USA) at 37 °C for 48 hours. The range of SMO concentrations was based on our recent study testing the efficacy of SMOs on total human PBMCs and malignant human B-cell lines^15^. Mature B-cell subsets were incubated with 333μg/mL of MTS (Promega, WI, USA) for 3 hours at 37 °C. Absorbance was measured at 490 nm with a TECAN Infinite M1000 Pro microplate reader.

### Analysis of bulk RNA-sequencing (RNA-seq) data

Raw sequencing reads were aligned to the GRCh38 and mm10 genome assembly for human and mouse, respectively, using the STAR package (v2.7.6a). Gene annotation, downloaded from the Ensembl database (v102), was used for STAR mapping and the subsequent read count evaluation. Picard (v2.26.11) was used to mark duplicates and generate the final bam files. HTSeq-count was employed to quantify the read counts for each gene, resulting in a count matrix. Then gene expression level was normalized by the variance stabilizing transformation (VST) algorithm in the DESeq2 package. Differential expression analyses were conducted using DESeq2.

### Analysis of scRNA-seq

For human and mouse scRNA-seq, the raw FASTQ file reads were aligned to the GRCh38 and mm10 genome assembly, respectively, using Cell Ranger v7.2.0 (10x Genomics). Cells were first filtered for quality control, removing those with fewer than 200 unique genes detected or greater than 10% mitochondrial reads. For mouse and human, doublets were identified and removed using DoubletFinder and scDblFinder R packages, respectively. The remaining high-quality cells were used to generate count matrices of gene expression levels per cell. Samples were then merged to create a Seurat object, joining data from individual samples. For mouse, each sample was normalized using SCTransform, selecting 3,000 features for integration. Principal component analysis (PCA) was computed for each object and integration was performed using reciprocal PCA in Seurat. FindIntegrationAnchors function was employed and IntegrateData function with dims = 50 was followed. After integration, PCA was recomputed on the integrated assay, and a UMAP plot was constructed using the first 50 principal components. For scRNA-seq of human samples, each sample was normalized using NormalizeData and FindVariableFeatures selecting 2,000 features for integration was followed. Sketch integration was performed sampling 10,000 representative cells from each sample utilizing a leverage score for each cell’s influence toward the gene-covariance matrix. FindVariableFeatures was performed again on the representative sketch object. Normalized gene counts of the sketch object were scaled using ScaleData and PCA was performed. For both mouse and human samples, cells were clustered using the FindNeighbors function with the first 50 principal components, followed by the FindClusters function with resolution = 1, after empirically evaluating multiple clustering resolutions. Finally, for human samples, extended cluster labels were extended to the full dataset of each sample by employing ProjectData. Gene expression was visualized by the FeauturePlot_scCustom (http://samuel-marsh.github.io/scCustomize) function using the default ‘order’ argument to superimpose positive cells on top in the UMAP plots. Differentially expressed genes (DEGs) for each cluster were identified using the FindAllMarkers function, applying the negative binomial test. For mouse samples, cell type annotation was manually conducted based on well-known marker genes of each cluster and later the clusters with the same cell type annotation were merged. For human samples, each cell was automatically annotated with scMRMA R package and validated by known major immune markers. Anti-and pro-apoptosis-related genes were selected based on relevant literature and pathway databases. The AUCell ^50^ package was then used to score each cell based on its expression of apoptosis-associated genes, reflecting the likelihood or intensity of apoptotic activity. The top 10 differentially expressed genes (**Fig. S19**) were identified for each B-cell subset in healthy mice. The AUCell package was then used to score each cell from healthy mice, control SMO-treated, and LFPRLR SMO-treated SLE-prone mice based on the top 10 genes of each B-cell subset derived from healthy mice.

### Gene set enrichment analysis (GSEA)

GSEA for comparing scRNA-seq data between any two groups was performed using Seurat’s FindMarkers between different subgroups generating a ranked differential gene expression list utilizing log_2_ fold change values and the Wilcoxon Rank Sum test. The ranked gene list was imported into either the Broad Institute’s GSEA tool or clusterProfiler R package to observe different pathways within the Gene Ontology (GO) gene sets.

### PRLR isoform annotation from bulk and scRNA-seq

The genomic coordinates of exon 10 of LF PRLR (human)/Prlr (mouse) were obtained from Ensembl. Similarly, coordinates were obtained for exon 11 of SF1a and SF1b PRLR in human, and exons 12, 11, and 13 of SF1, SF2, and SF3 Prlr, respectively, in mice. Reads, mapped to specific exons noted above, were extracted using samtools from bulk and scRNA-seq data. These reads were subsequently utilized to represent the expression of the long and short isoforms of PRLR. The number of unique molecular identifiers (UMIs) of each exon for each cell was obtained for further single-cell analysis.

### Immunoglobulin sequencing

Raw FASTQ files were processed with the Cell Ranger VDJ pipeline (v7.2.0), which aligned reads to the appropriate reference genome and annotated V(D)J rearrangements. Cells with high confidence, productive complementary determining 3 (CDR3) region calls were retained. For each retained cell, the single CDR3 sequence supported by the greatest number of UMIs was selected. The length of the peptides of CDR3 was checked between control SMO and LFPRLR SMO to see if there were any changes characteristic of autoimmunity^55^. CDR3s were classified as long [>15 amino acids (aa)] or short (<15 aa). The amino acid composition of every CDR3 was expressed as residue proportions, log transformed, and visualized with the ComplexHeatmap R package.

### Statistics

Unless otherwise noted, statistical analysis was performed using unpaired two-tailed Student’s t-and Mann-Whitney U-tests (for comparisons of two groups) or one-way ANOVA (for comparison of three or more groups) with Dunnett’s multiple-comparisons test, using GraphPad Prism (version 9.5.1; GraphPad Software, San Diego, CA, USA). Correlation analyses were performed using the Spearman method, and two-tailed p-values were calculated using a simple linear regression t-test. P-values for proportion analyses for mouse scRNA-seq data were calculated using a chi-square test. Female SLE-prone mice and samples from female patients with SLE were utilized because the incidence of SLE in females is 9-fold that in males. The number of samples utilized in each experiment was calculated using the G*power function and based on the variation within each group known from our previous studies ^15^. Results are expressed as mean ± s.e.m.; p < 0.05 was considered statistically significant, 0.05 ≤ p ≤ 0.1 was considered a trend.

## Data availability

The three normal mouse single-cell data were obtained from the 10X genomics public database: https://www.10xgenomics.com/datasets/mouse-splenocytes-5-v-2-whole-transcriptome-analysis-2-standard-6-0-1, https://www.10xgenomics.com/datasets/10k-Mouse-Splenocytes-5p-gemx and https://www.10xgenomics.com/datasets/5k-Mouse-Splenocytes-5p-nextgem. The scRNA-seq data for splenic leukocytes of SLE-prone *MRL-lpr* mice subjected to treatment with control SMO or LFPRLR SMO were deposited in Gene Expression Omnibus (GEO) under GSE240076. Publicly available GEO datasets GSE137029, GSE211700, and GSE186367 were analyzed.

## Code and software

Code is available upon request. All software packages used for analysis are provided in **Table S6**.

## Abbreviations

aa: amino acid
APRIL: A proliferation-inducing ligand
BAFF: B-cell activating factor
BCR: B-cell receptor
CD: cluster of differentiation
CDR3: complementary determining region 3
CM: classical monocytes
DAPI: 4′,6-diamidino-2-phenylindole
DC: dendritic cell
DMSO: dimethyl sulfoxide
DNA: deoxyribonucleic acid
DSMZ: Deutsche Sammlung von Mikroorganismen und Zellkulturen
EdU: 5-ethynyl-2′-deoxyuridine
FAS: fas cell surface death receptor
FBS: fetal bovine serum
FPKM: fragments per kilobase of transcript per million mapped reads
GC: germinal center
GSEA: gene set enrichment analysis
IACUC: Institutional Animal Care and Use Committee
IC50: half-maximal inhibitory concentration
IFN-I: type I interferon
IFPRLR: intermediate isoform of PRLR
IGH: immunoglobulin heavy chain
JAK: Janus kinase
JAX: Jackson Laboratories
LFPRLR: long isoform of PRLR
MACS: magnetic activated cell sorting
MFI: median fluorescence intensity
mRNA: messenger ribonucleic acid
MRL/lpr: Murphy Roths Large lymphoproliferation
MTS: (3-(4,5-dimethylthiazol-2-yl)-5-(3-carboxymethoxyphenyl)-2-(4-sulfophenyl)-2H-tetrazolium)
I/NCM: intermediate and non-classical monocytes
NGS: next generation sequencing
NK: natural killer
PBMC: peripheral blood mononuclear cells
PRL: prolactin
PRLR: prolactin receptor
RNA-seq: RNA sequencing
scRNA-seq: single cell RNA sequencing
SFPRLR: short isoform of PRLR
SLE: systemic lupus erythematosus
SMO: splice modulating oligomer
STAT: signal transducer and activator of transcription
UB: ubiquitin
WBC: white blood cell
UMI: unique molecular identifiers
NES: normalized enrichment score
siRNA: small interfering RNA.

## Author Contributions

Conceptualization: SS, AMW

Methodology: SS, AMW, KH, ZH, JRL-J, IS, EM, JLK, XW, ZG, KM

Investigation: KH, ZH, JRL-J, ATK, AK, ASO, XG, DHJ, HQ, ST, MYL

Visualization: SS, AMW, KH, ZH, JRL-J Funding acquisition: SS, AMW

Project administration: SS, AMW Supervision: SS

Writing - Original Draft: SS

Writing – review & editing: AMW, KH, ZH, JRL-J, ATK, AK, HQ, ST, IS, EM, JLK, XW, ZG, KM

## Acknowledgements

We thank Dr. Nora Heisterkamp (City of Hope) for productive discussions.

Schematics were created using BioRender.com.

This work was supported by the following grants:

R37CA276517-01A1 from the National Cancer Institute of the National Institutes of Health (SS)

R21AR084116-01 from the National Institutes of Arthritis, Musculoskeletal and Skin Diseases and the Office of the Director of the National Institutes of Health (SS)

American Society of Hematology Bridge Grant (SS) American Society of Hematology Scholar Award (SS)

University of California Drug Discovery Consortium and Ono Pharmaceuticals (AMW) Conquer Cancer Now Award from the Concern Foundation (SS)

Congressional Families Program Award from the Prevent Cancer Foundation (SS) Shared Resources Pilot Grant from City of Hope (SS)

City of Hope-University of California Riverside Biomedical Research Initiative Award (SS, AMW)

Analytical Cytometry Core and Integrative Genomics Core Shared Resources are supported by the National Cancer Institute of the National Institutes of Health under grant number P30CA033572. The content is solely the responsibility of the authors and does not necessarily represent the official views of the National Institutes of Health.

## Data and materials availability

GSE numbers for all single-cell and bulk RNA sequencing data used in this study are provided in Materials and Methods. No new code was generated. Sequences of control and LFPRLR SMO are published ^15^. SMOs are covered by a pending US patent application and will be shared as part of material transfer agreement between City of Hope/University of California Riverside and the recipient institution.

