## Supplemental figures and tables for "Knockdown of the long isoform of the prolactin receptor selectively targets pathogenic immune cells in systemic lupus erythematosus and averts glomerular pathology"

### Inventory

- Supplementary Figures S1-S20
- Supplementary Tables S1-S5

*Fig S1. Steps for the analysis of scRNA-seq of PBMCs from healthy donors (n = 12) and patients with SLE (n = 34) from GSE137029.*

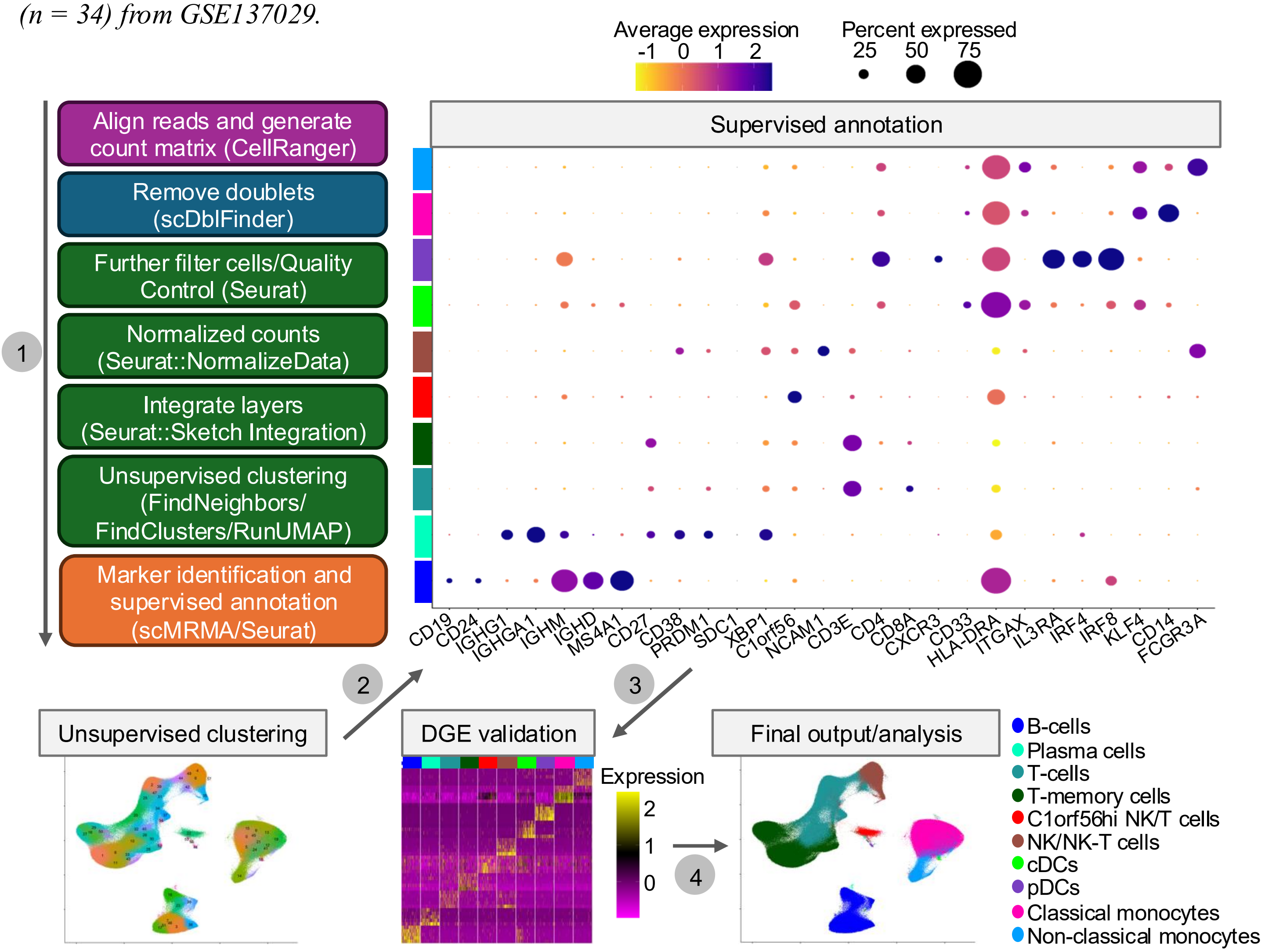


**Fig S1. Steps for the analysis of scRNA-seq of PBMCs from healthy donors (n = 12) and patients with SLE (n = 34) from GSE137029.**

***Fig S2.*** *Monocytes, the highest PRLR-expressing human PBMC subset, are aberrantly enhanced at the expense of lymphoid cell subsets within the PRLR⁺ fraction in SLE patients*

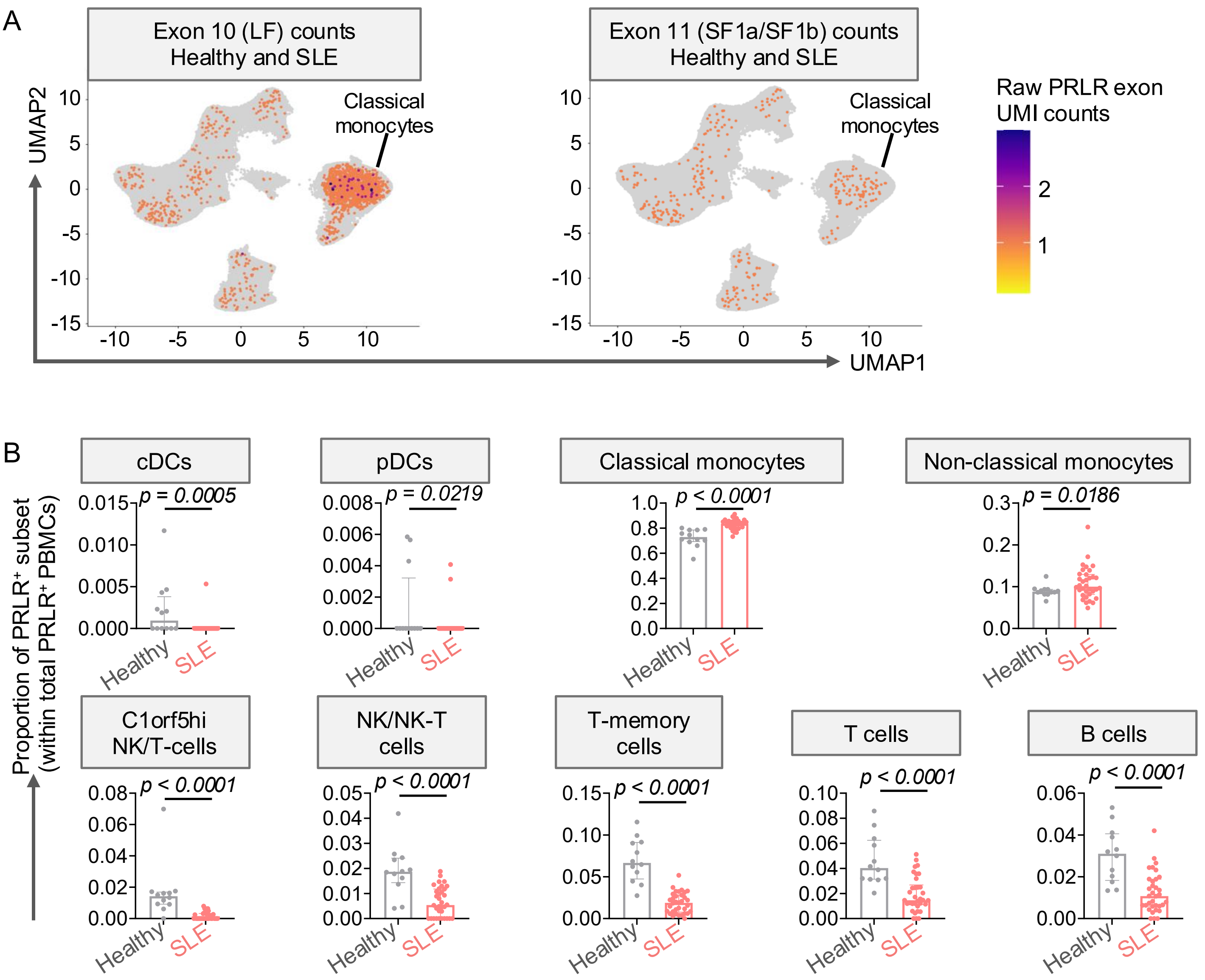


**Fig S2. Monocytes, the highest PRLR-expressing human PBMC subset, are aberrantly enhanced at the expense of lymphoid cell subsets within the PRLR⁺ fraction in SLE patients.** (**A**) UMAP plots depicting total UMI counts of exon 10 (LFPRLR), exon 11 (SF1a/b PRLR) in heathy donors (n=12) and patients with SLE (n=34). (**B**) Proportions of each immune cluster within the PRLR⁺ fraction were calculated and compared between healthy donors (n=12) and patients with SLE (n=34). Bar charts: median ± interquartile range, p-values: Mann-Whitney U test.

***Fig S3.*** *LFPRLR is aberrantly upregulated in immune cells but not in kidney and brain in SLE-prone mice*

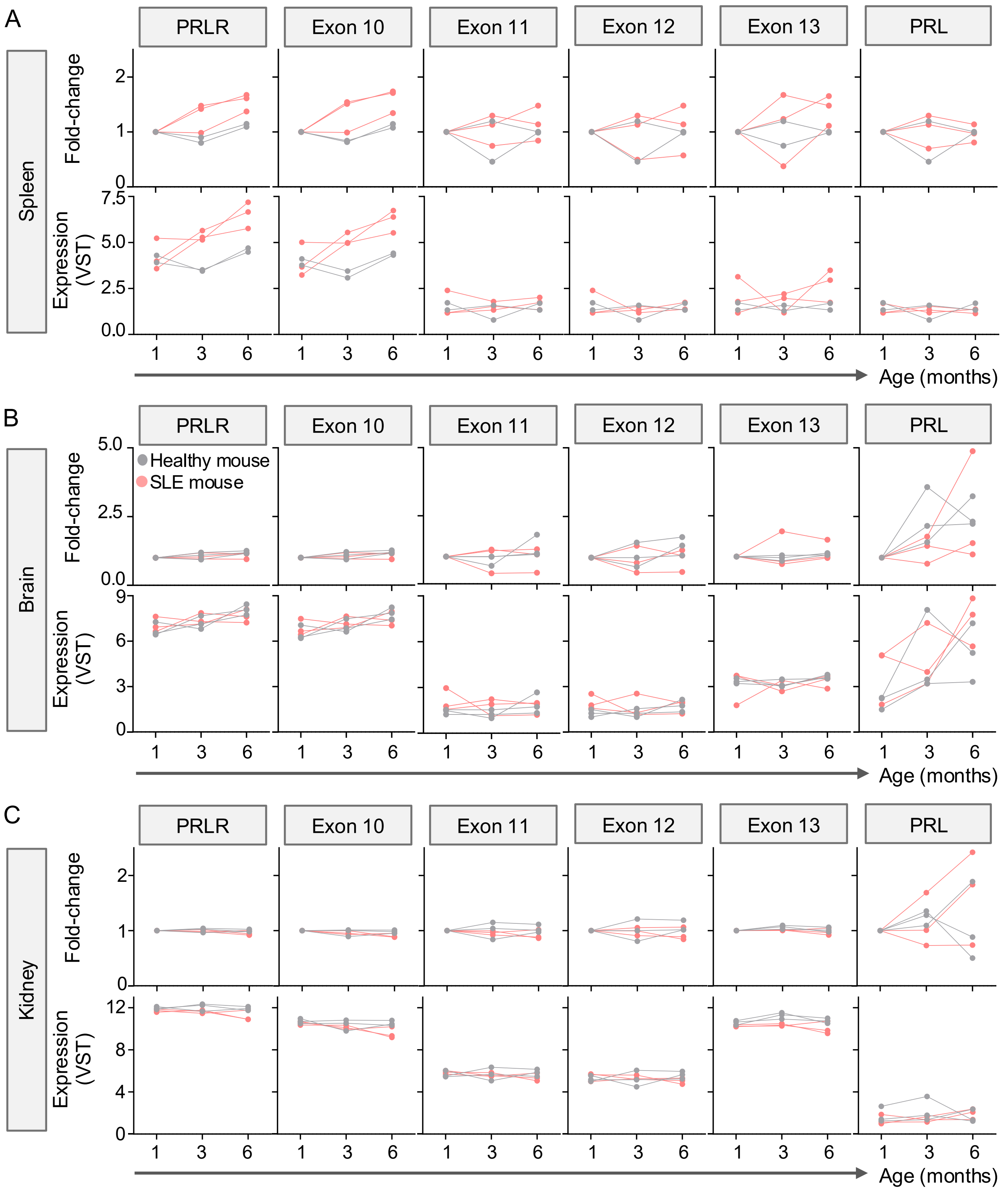


**Fig S3. LFPRLR is aberrantly upregulated in immune cells but not in kidney and brain in SLE-prone mice.** Total PRLR, LFPRLR (exon 10), SF2 PRLR (exon 11), SF1 PRLR (exon 12), SF3 PRLR (exon 13), and PRL transcript expression by bulk RNA-seq (GSE186367) in (**A**) spleen, (**B**) brain, and (**C**) kidney of 1-month-, 3-month- and 6-month-old *NZB/W-F1* SLE-prone mice (n=3) and age- and sex-matched healthy mice (n=3). Fold-change of expression in each gene was calculated with respect to the expression measured at 1 month of age.

***Fig S4.*** *LFPRLR SMO changes PRLR isoform expression by reducing the LF and/or increasing the SFs in healthy donor and SLE patient PBMCs*

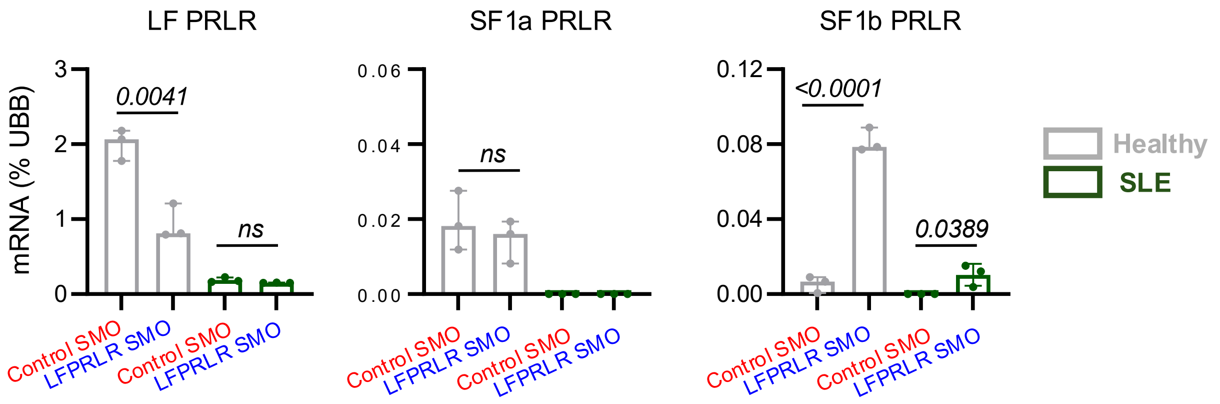


**Fig. S4. LFPRLR SMO changes PRLR isoform expression by reducing the LF and/or increasing the SFs in healthy donor and SLE patient PBMCs.** Transcript levels of PRLR isoforms LF, SF1a, and SF1b quantified by qPCR in healthy donor (n=3) or SLE patient PBMCs (n=3) treated *ex vivo* with control SMO or LFPRLR SMO for 24h. Transcripts were normalized to UBB and expressed as % of UBB. Each dot represents an independent technical replicate (healthy donor, grey; SLE patient, green). One representative of three biological replicates is shown. Median ± interquartile range, p-values: unpaired two-tailed Student's t test. ns = not significant

*Fig S5.* *Gating strategy for high-dimensional flow cytometry analysis of human PBMCs*

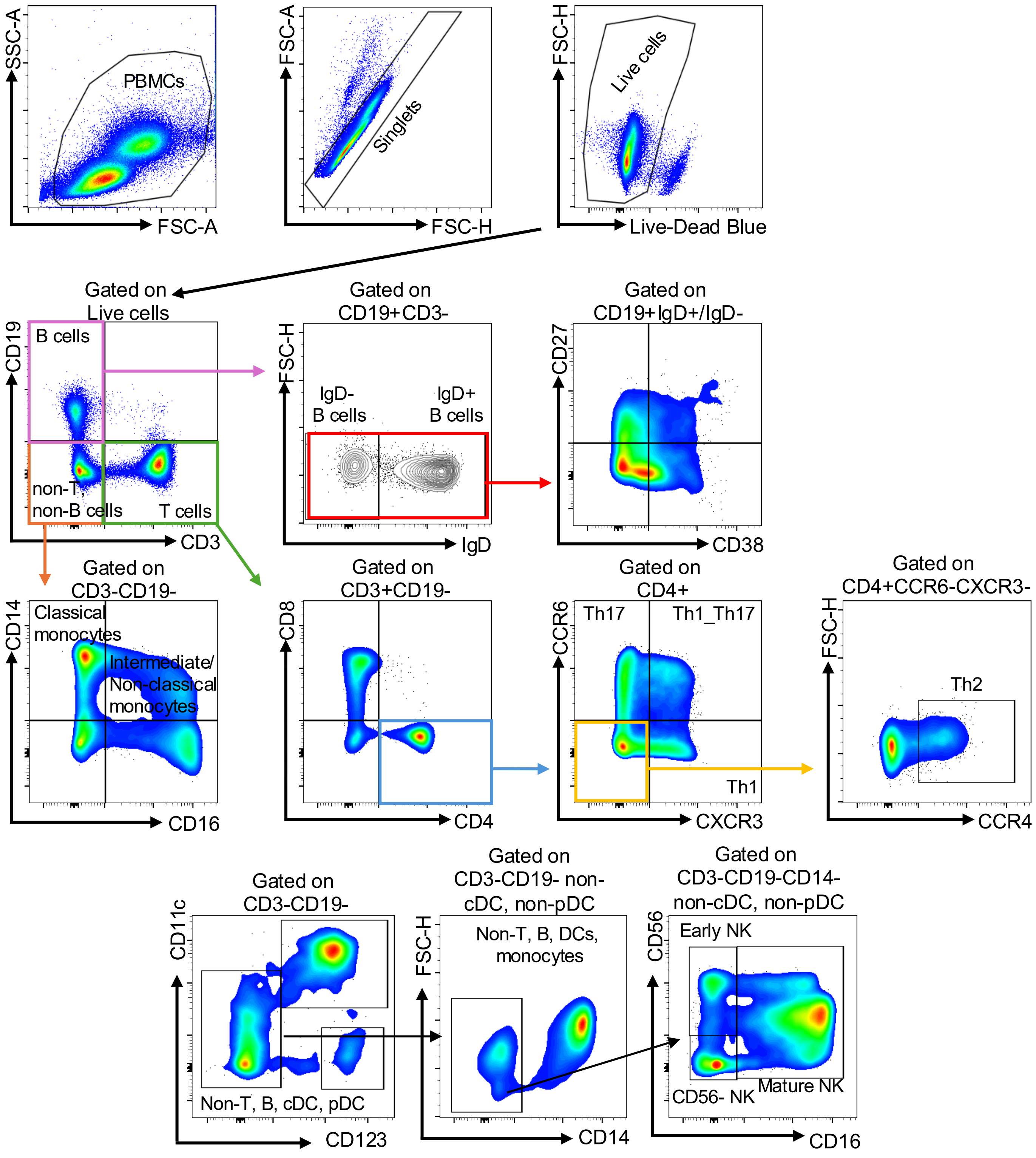


**Fig S5. Gating strategy for high-dimensional flow cytometry analysis of human PBMCs.** Major immune lineages were measured in live, singlet fraction. B cells were gated as $CD19^{+}$ and subdivided into $IgD^{+}$ and $IgD^{-}$ subsets, followed by further stratification into CD27 and CD38 expression. T helper cells were gated as $CD3^{+}CD4^{+}$ and then classified into Th1 ($CD4^{+}CXCR3^{+}CCR6^{-}$), Th17 ($CD4^{+}CXCR3^{-}CCR6^{+}$), Th1/Th17 ($CD4^{+}CXCR3^{+}CCR6^{+}$), and Th2 ($CD4^{+}CXCR3^{-}CCR6^{-}CCR4^{+}$) subsets. Monocytes within $CD3^{-}CD19^{-}$ cells were identified as classical ($CD14^{+}CD16^{-}$) and intermediate/non-classical ($CD14^{+}CD16^{+}$) subsets. NK cells were gated as non-T, non-B, non-monocyte, non-cDCs, non-pDCs, and classified into early NK ($CD56^{bright}CD16^{-}$) and mature NK ($CD56^{dim}CD16^{+}$).

***Fig S6. LFPRLR knockdown for 72h ex vivo does not affect viability of healthy donor and SLE patient PBMCs***

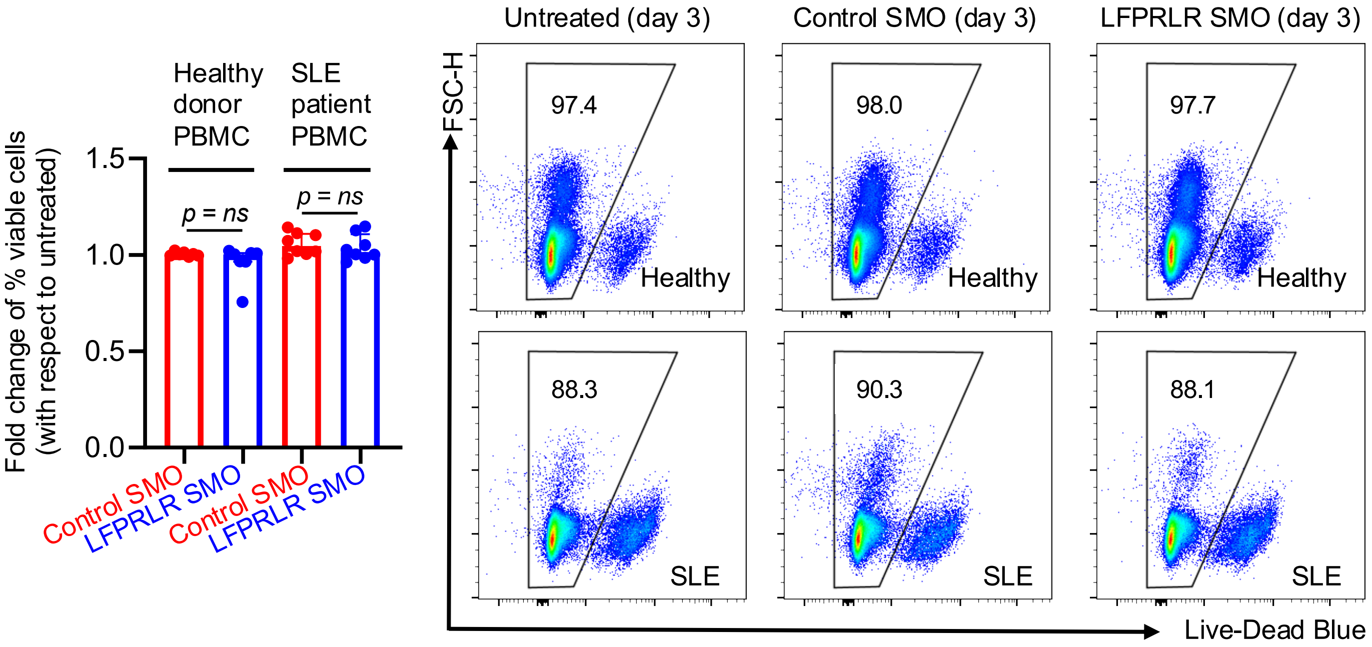


**Fig S6. LFPRLR knockdown for 72h *ex vivo* does not affect viability of healthy donor and SLE patient PBMCs.** Representative flow cytometry plots showing percentages of live PBMCs from SLE patients (n=8) and healthy donors (n=7) after 3 days of culture under three conditions: untreated, control SMO, and LFPRLR SMO. Live cells were gated as singlets and Live/Dead Blue staining. Data are represented as median ± interquartile range. P-value: not significant, calculated by Mann-Whitney U test. ns = not significant.

**Fig S7.** *Schema and validation of generation of major mature B-cell subsets from human PBMCs*

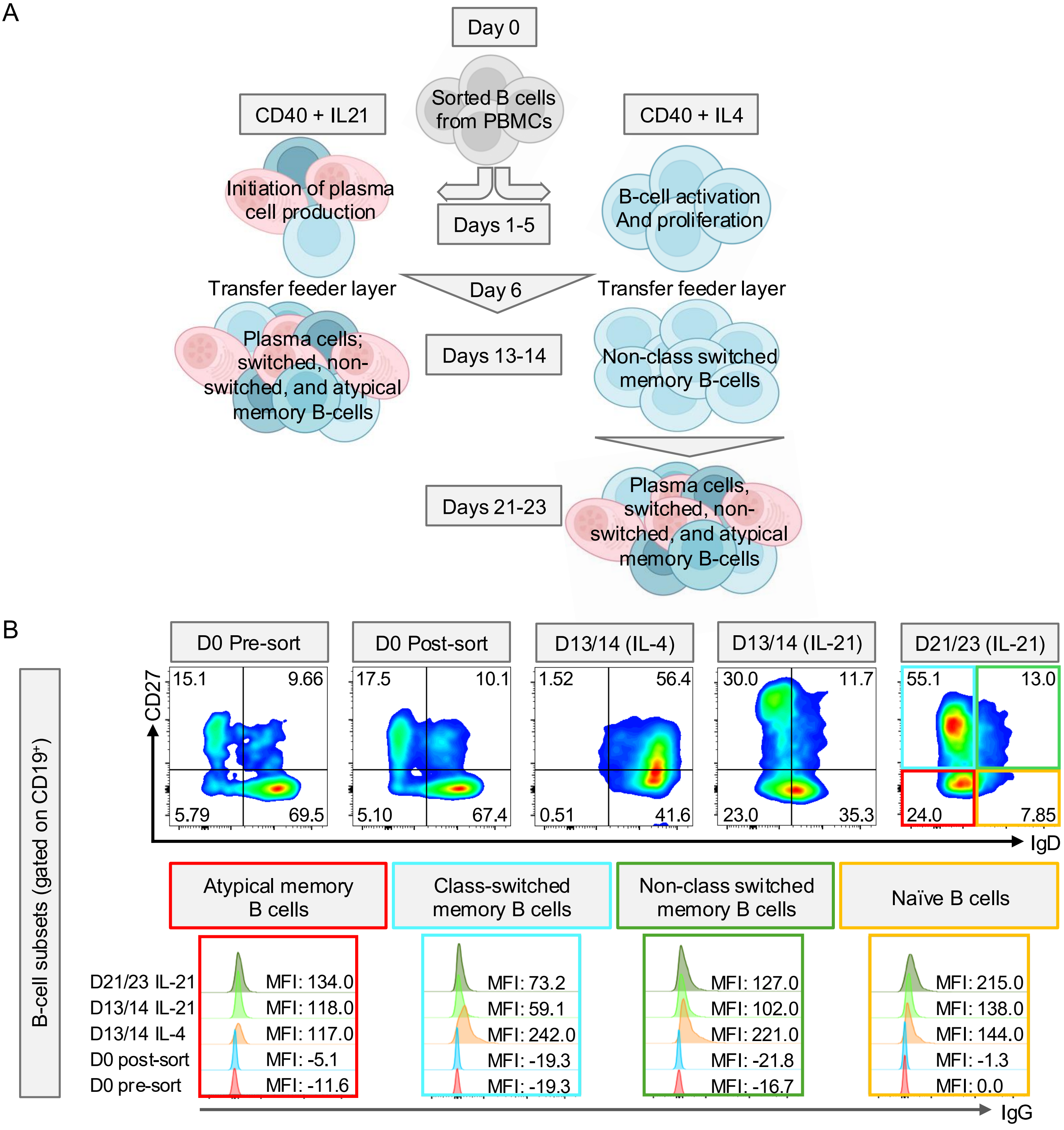


**Fig S7. Schema and validation of generation of major mature B-cell subsets from human PBMCs.** (**A**) In-house protocol for differentiating all major mature B-cell subsets from healthy donor PBMCs. B cells isolated from healthy donor PBMCs were cultured with IL-4 or IL-21 to generate distinct B-cell subsets. (**B**) Generation of mature B-cell subsets (activated and memory) from naïve B cells of healthy donors (n=8) confirmed by flow cytometry.

**Fig S7 contd.** *Schema and validation of generation of major mature B-cell subsets from human PBMCs*

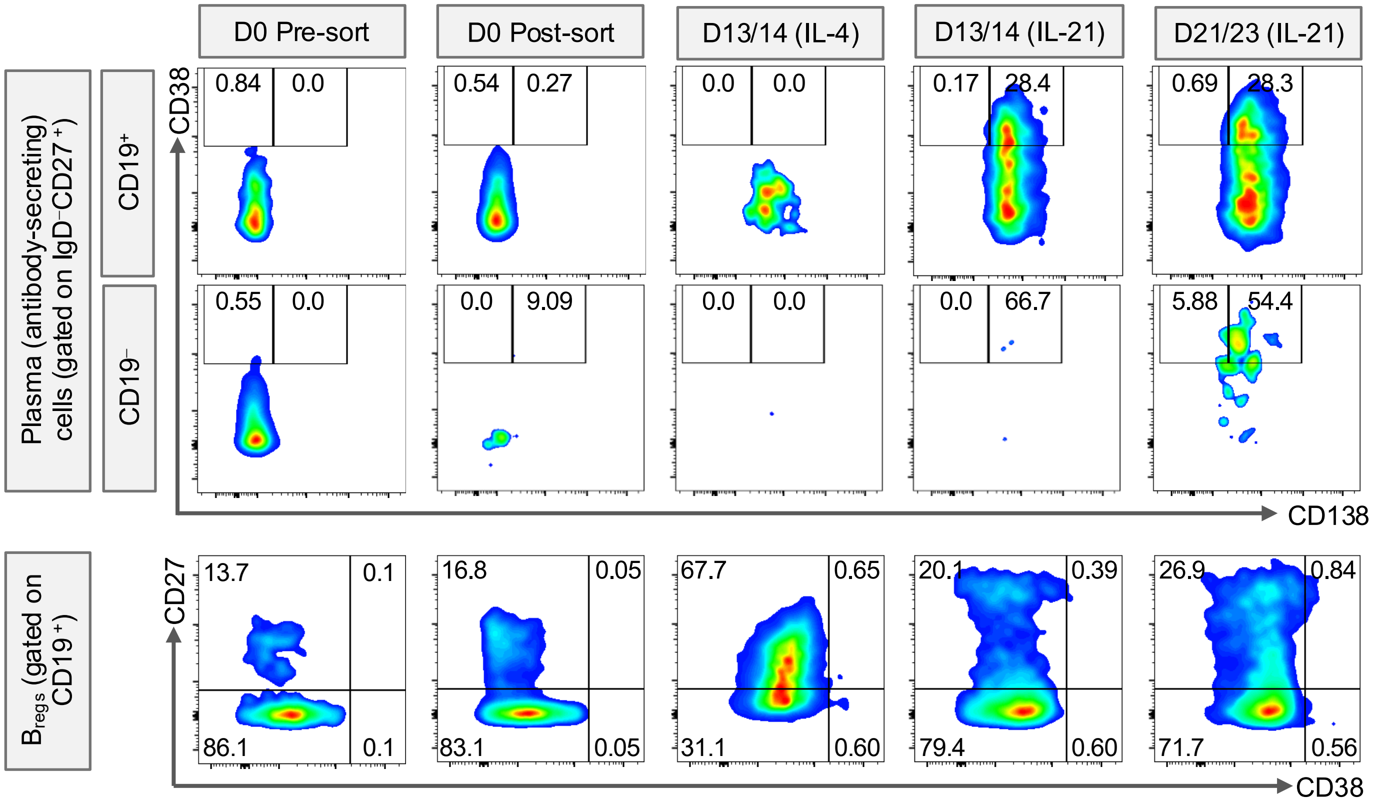


**Fig S7. Schema and validation of generation of major mature B-cell subsets from human PBMCs.** (**C**) Generation of mature B-cell subsets (plasma cells, and B-regulatory cells) from naïve B cells of healthy donors (n=8) confirmed by flow cytometry.

**Fig S8.** *Ex vivo knockdown of the LFPRLR does not impact viability or phenotype of normal humoral immune cells*

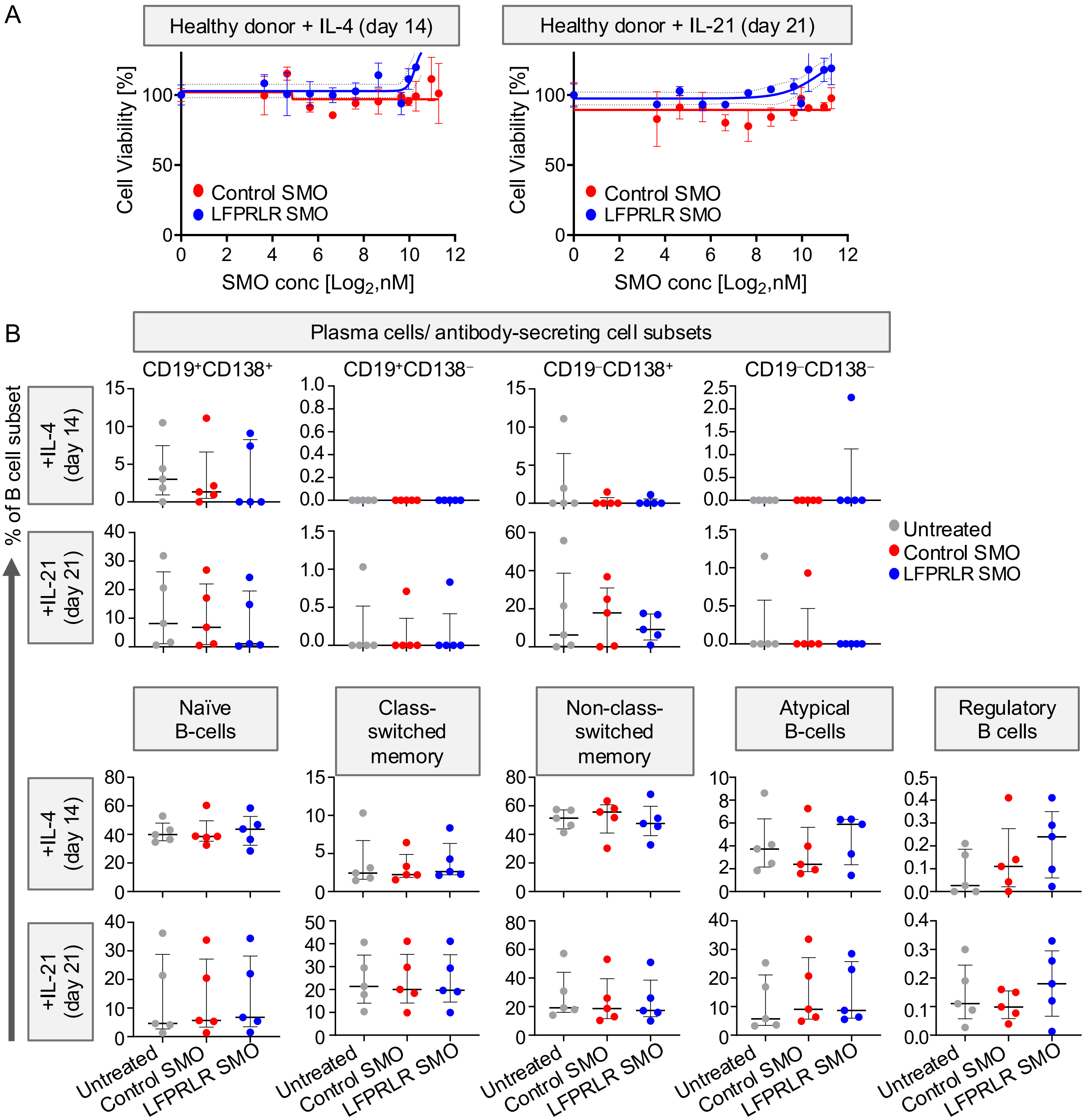


**Fig. S8. *In vitro* knockdown of the LFPRLR does not impact viability or phenotype of normal humoral immune cells.** (**A**) MTS assay assessing the viability of mature B-cell subsets in healthy donor PBMCs after treatment with control SMO or LFPRLR SMO for 48 h. Data from one of 9 healthy donors is shown. Data are represented as mean ±SEM. P-values: not significant, calculated by Student's t-test. (**B**) Frequency of plasma cell subsets, $IgD^{-}CD27^{+}$ class-switched memory B cells, $IgD^{-}CD27^{-}$ atypical memory B cells, $IgD^{+}CD27^{+}$ non-class-switched memory B cells, and $CD38^{bright}CD27^{-}$ regulatory B cells in cultures treated with control or LFPRLR SMO at day 14 (+IL-4) or day 21 (+IL-21) (n = 5). Data are represented as median ± interquartile range. P-values: not significant, calculated by Mann-Whitney U test.

**Fig S9.** *Steps for the analysis of scRNA-seq of mouse splenic immune cells*

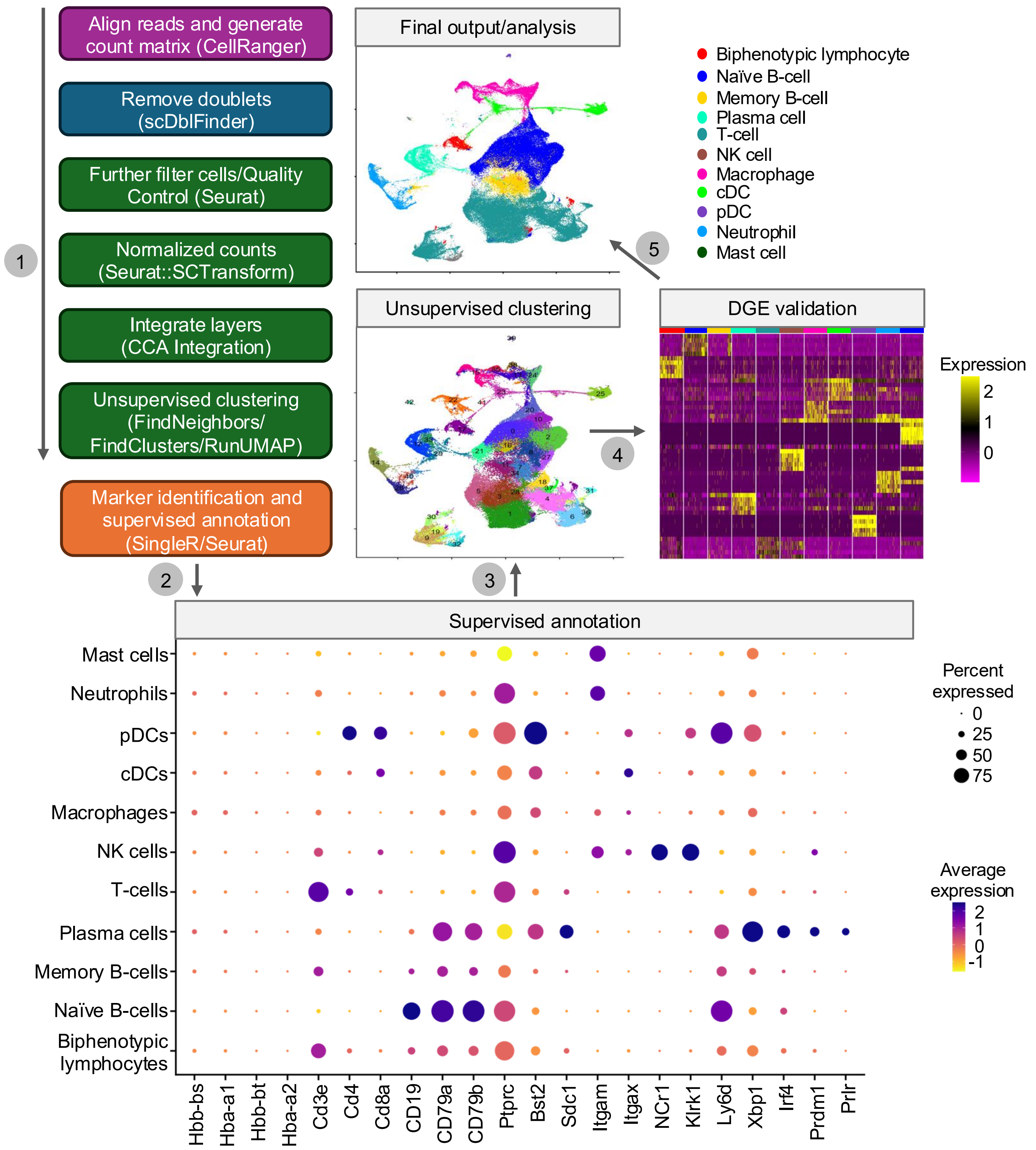


**Fig S9. Steps for the analysis of scRNA-seq of mouse splenic immune cells.** scRNA-seq workflow (steps 1-5) for analysis of splenic immune cells from healthy *C57BL/6J* (n=3), control SMO-treated *MRL-lpr* (n = 4), and LFPRLR SMO-treated *MRL-lpr* (n=4) mice.

***Fig S10.*** *LFPRLR is the predominant PRLR isoform and is most expressed in splenic plasma cells of normal and SLE-prone mice*

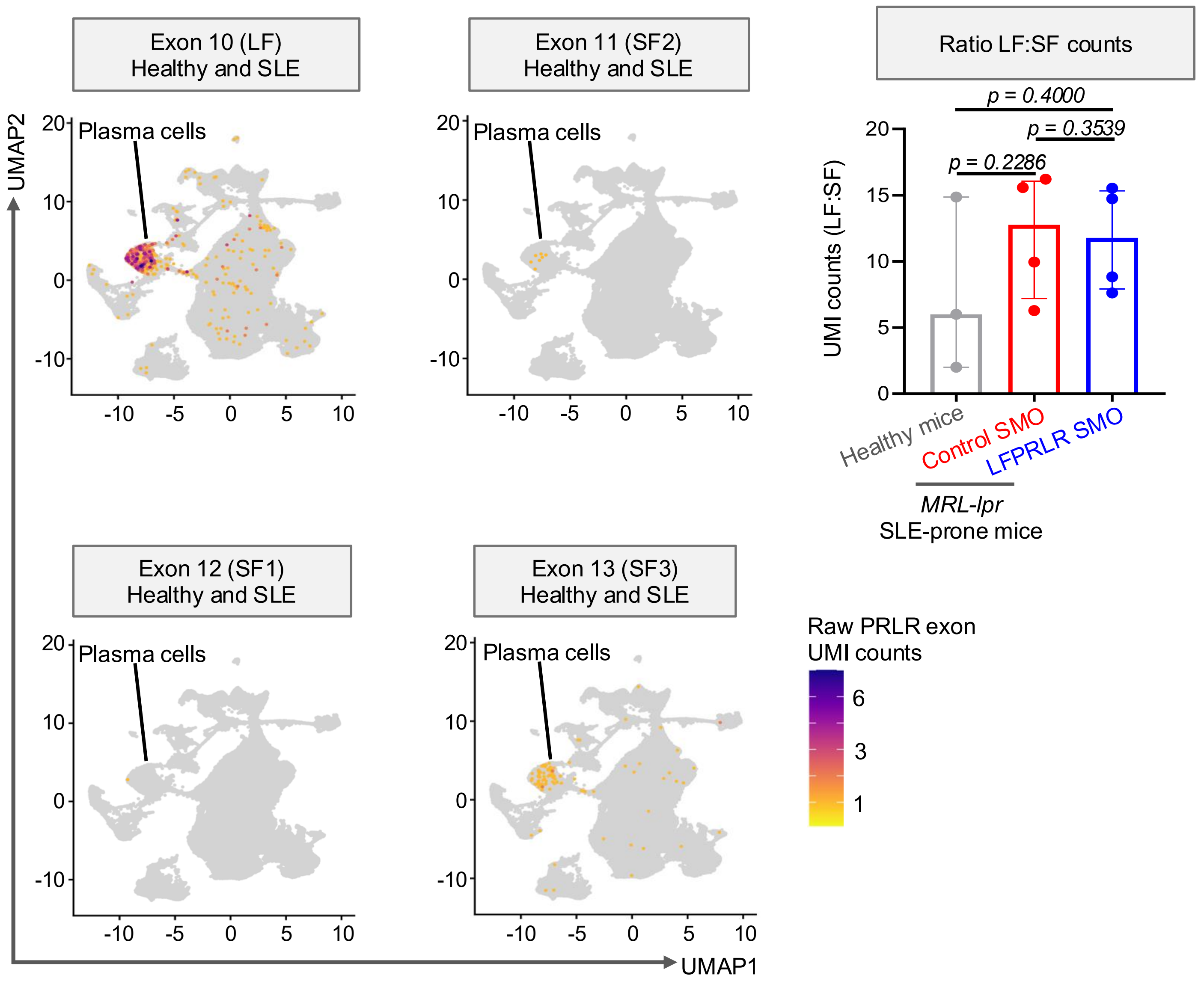


**Fig S10. LFPRLR is the predominant PRLR isoform and is most expressed in splenic plasma cells of normal and SLE-prone mice.** UMI counts of exon 10 (LFPRLR), exon 11 (SF1 PRLR), exon 12 (SF3 PRLR), exon 13 (SF2 PRLR) were extracted from healthy *C57BL/6J* (n=3), control SMO-treated *MRL-lpr* (n=4), and LFPRLR SMO-treated *MRL-lpr* (n=4) mice. The ratio of UMI counts between LF:SF in each mouse was calculated. Bar charts: median ± interquartile range, p-values: Mann-Whitney U test.

*Fig S11. Plasma cells, macrophages and neutrophils are aberrantly enhanced at the expense of B- and T-cell subsets in the PRLR⁺ fraction in SLE mice*

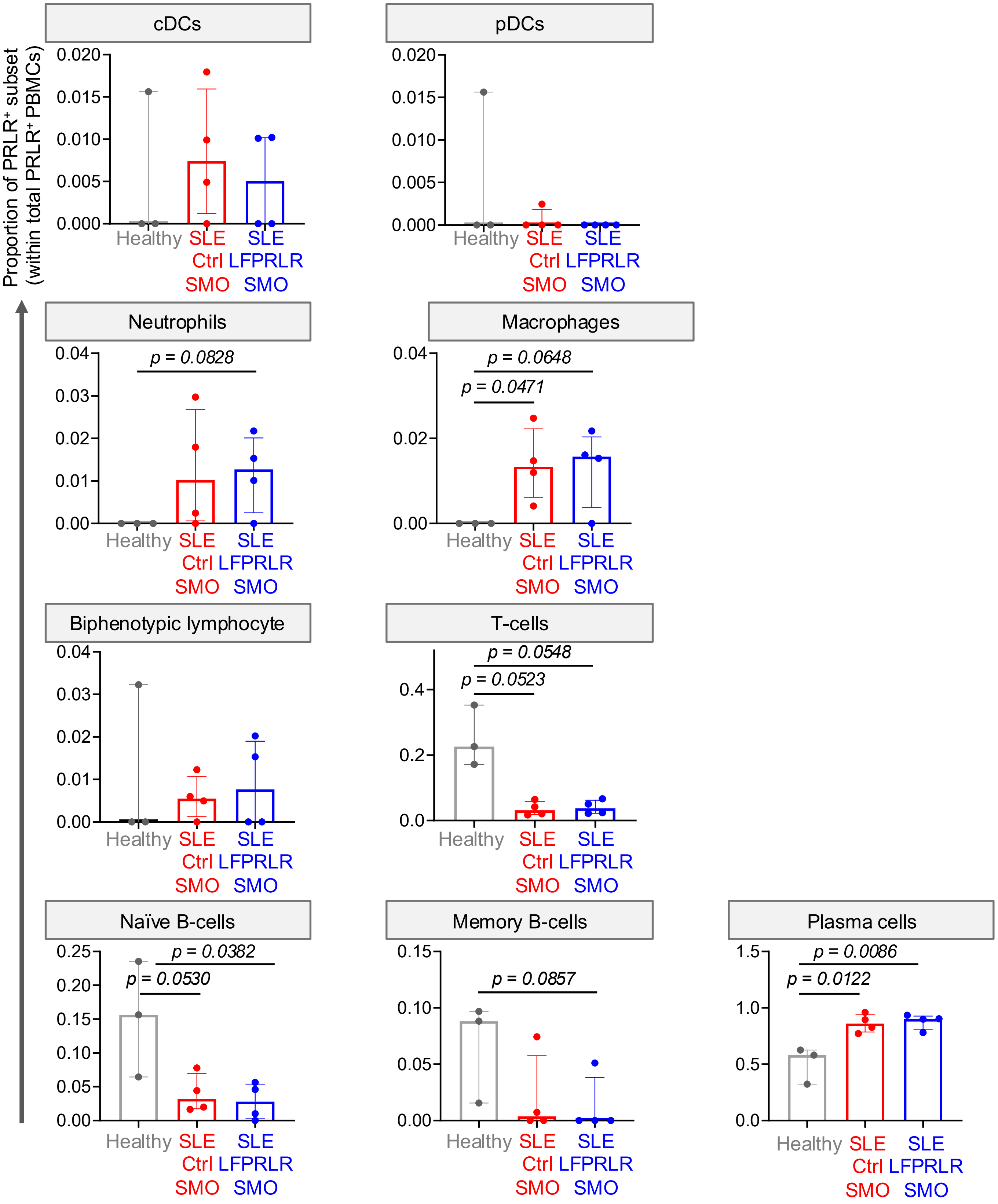


**Fig S11. Plasma cells and macrophages are aberrantly enhanced at the expense of B- and T-cell subsets in the PRLR⁺ fraction in SLE mice.** Proportions of each immune cluster within the PRLR+ fraction were compared between healthy (n=3), LFPRLR SMO-treated (n=4), and control SMO-treated mice (n=4). Bar charts: median ± interquartile range, p-values: Mann-Whitney U test.

***Fig S12.*** *LRPRLR knockdown reduces the absolute numbers of STAT1/2/5⁺ PRLR⁺ plasma cells and STAT1⁺ PRLR⁺ macrophages in spleens of SLE-prone mice.*

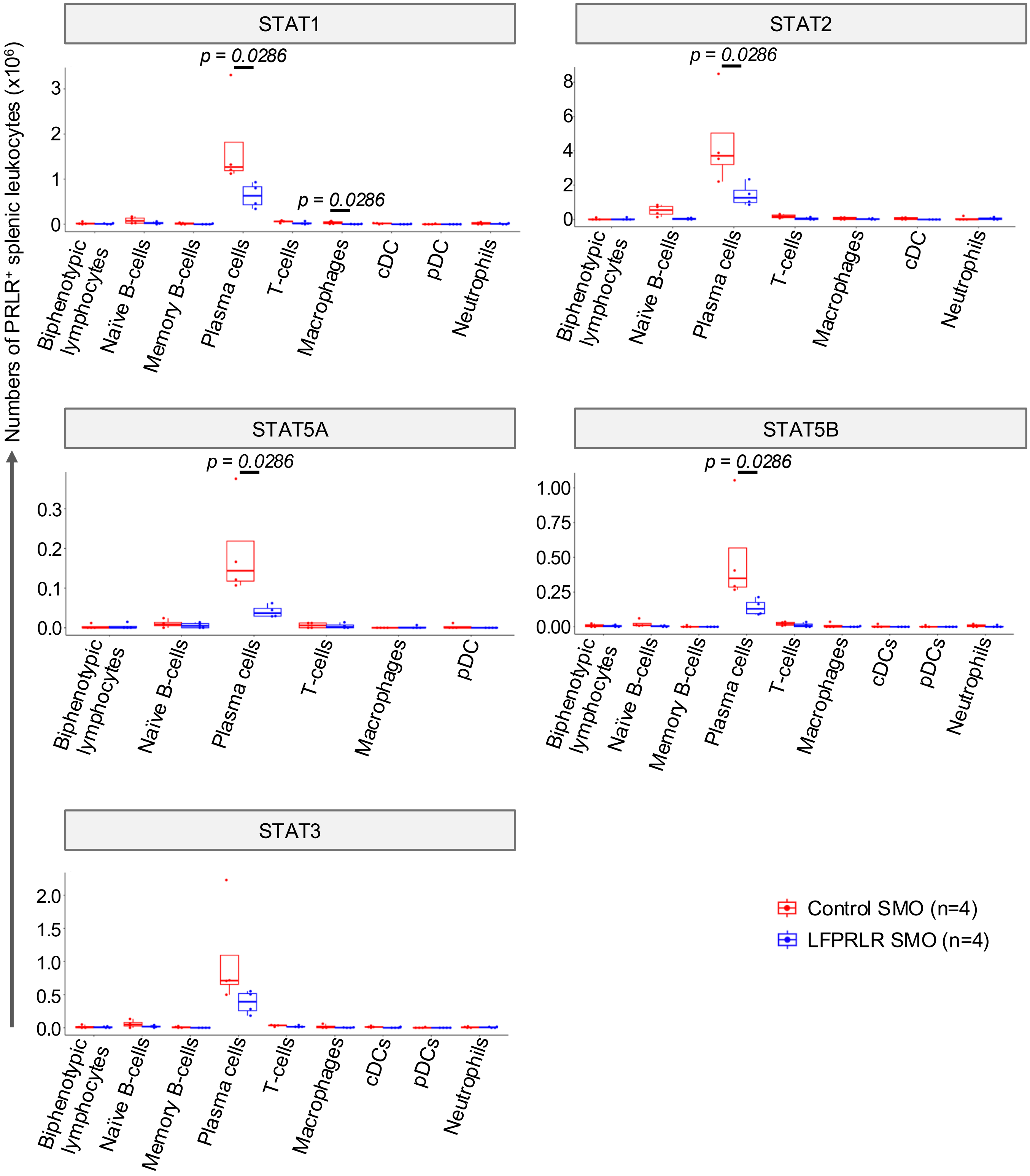


**Fig S12. LRPRLR knockdown reduces the absolute numbers of STAT1/2/5⁺ PRLR⁺ plasma cells and STAT1⁺ PRLR⁺ macrophages in spleens of SLE-prone mice.** Absolute counts of PRLR⁺ STAT⁺ splenic leukocyte subsets in *MRL-lpr* mouse treated with either control SMO (n=4) or LFPRLR SMO (n=4). Box plots: median ± interquartile range, p-values: Mann-Whitney U test. Only significant p-values are shown.

***Fig S13.*** *LRPRLR knockdown reduces the absolute numbers of STAT1/5⁺ PRLR⁻ plasma cells.*

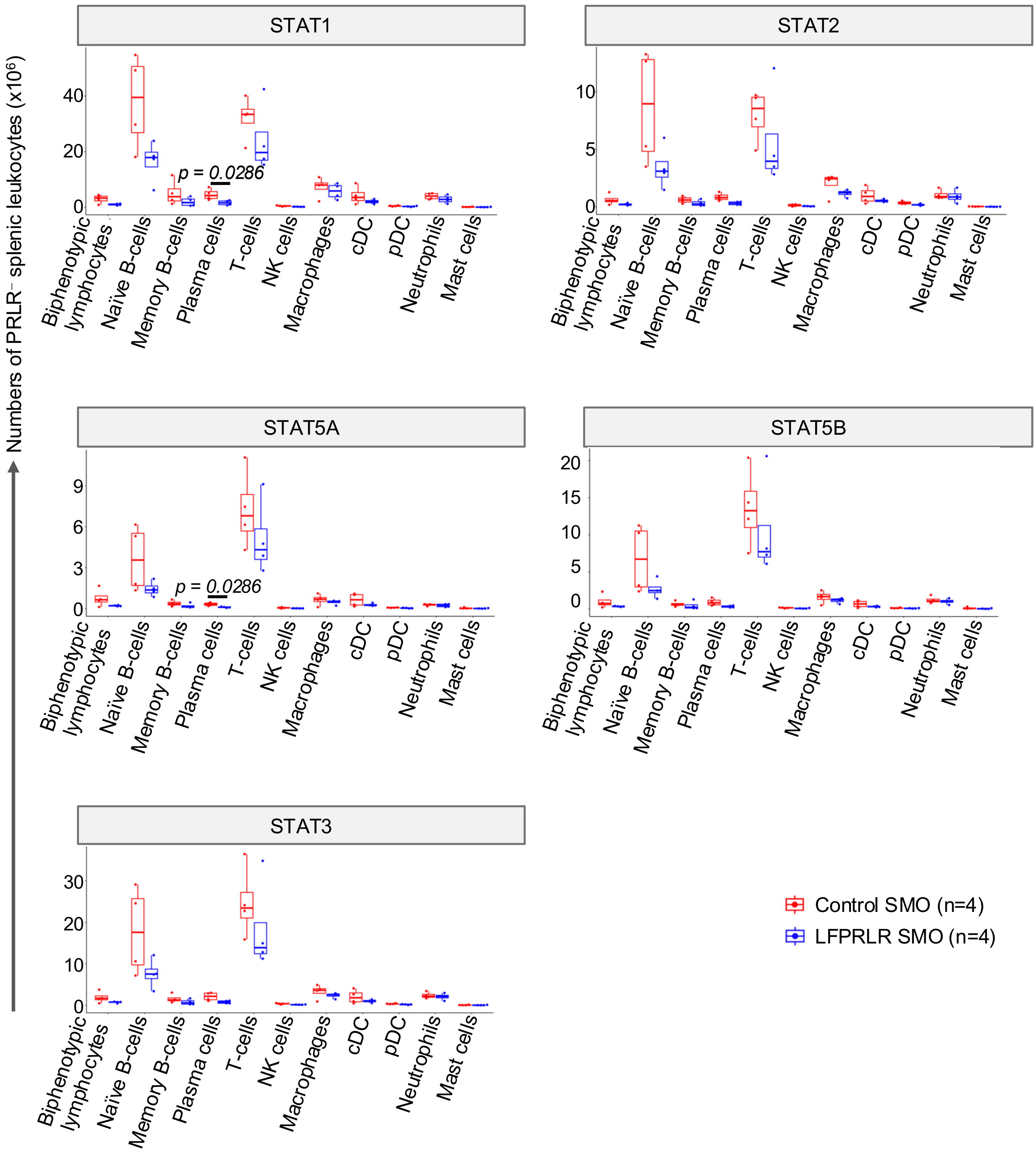


**Fig S13. LRPRLR knockdown reduces the absolute numbers of STAT1/5⁺ PRLR⁻ plasma cells.** Absolute counts of PRLR⁻ STAT⁺ splenic leukocyte subsets in *MRL-lpr* mouse treated with either control SMO (n=4) or LFPRLR SMO (n=4). Box plots: median ± interquartile range, p-values: Mann-Whitney U test. Only significant p-values are shown.

*Fig S14. Downstream Tlr4 signaling molecules involved in activation and subsequent maturation into plasma cells were reduced in B-cell subsets of SLE-prone mice after LFPRLR knockdown*

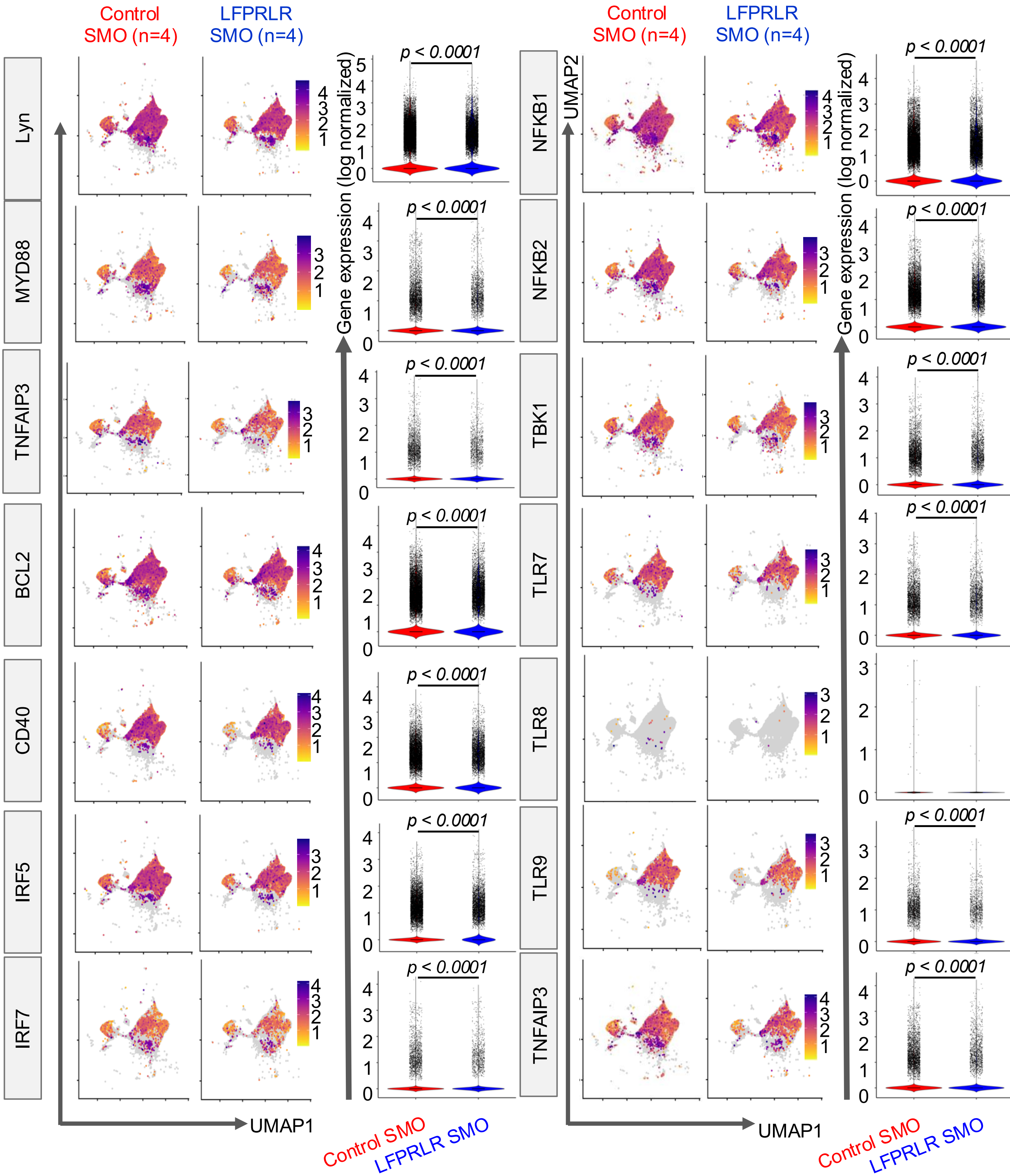


**Fig S14. Downstream Tlr4 signaling molecules involved in activation and subsequent maturation into plasma cells were reduced in B-cell subsets of SLE-prone mice after LFPRLR knockdown.** UMAP plots and gene expression distributions Tlr4 signaling transcripts in *MRL-lpr* splenic B cells after LFPRLR knockdown of LFPRLR. Violin plots: Log-normalized counts, p-values: Mann-Whitney U test.

*Fig S15.* *LFPRLR knockdown in SLE-prone mice has no effect on apoptosis pathway genes in splenic immune cells*

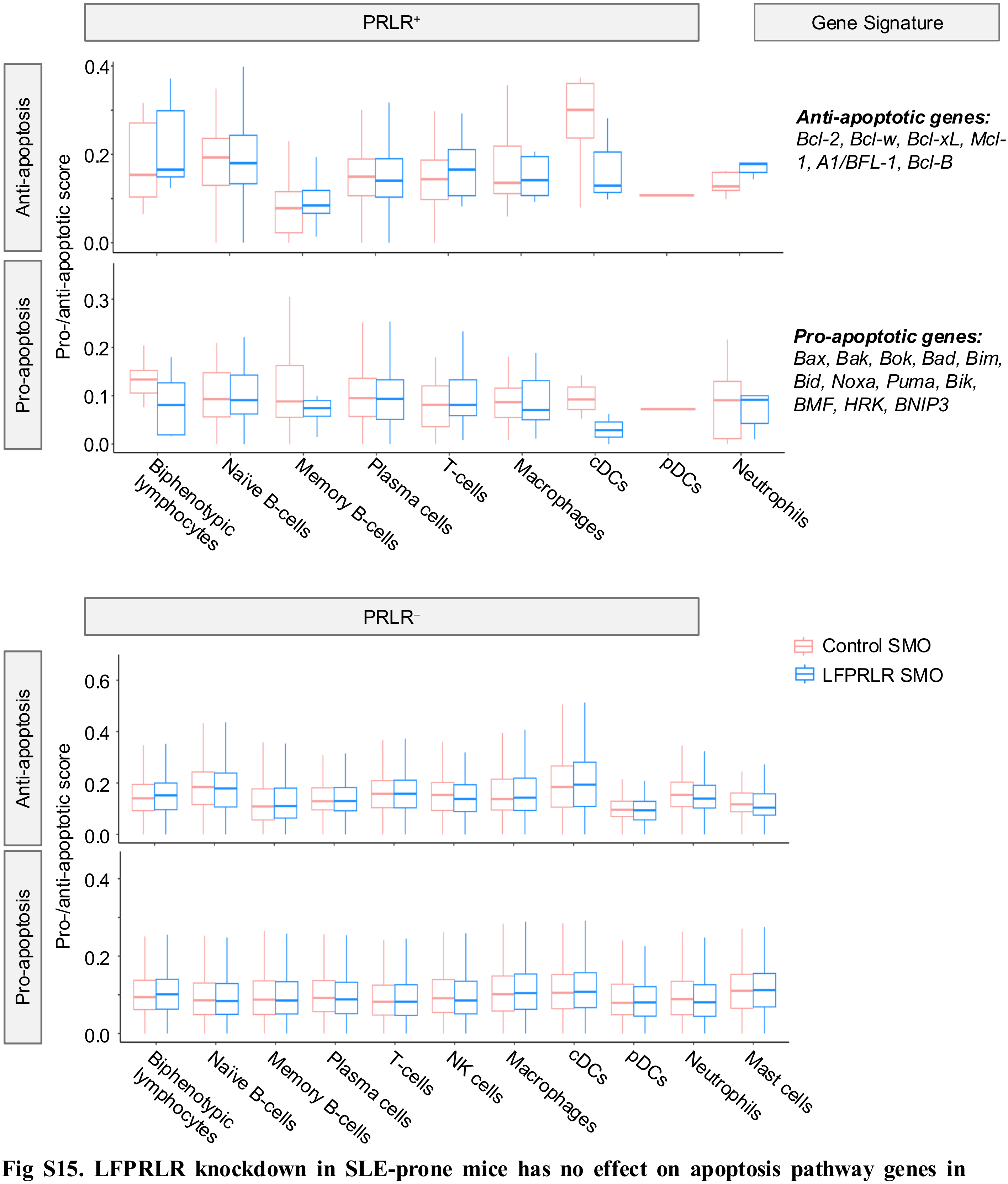


**Fig S15. LFPRLR knockdown in SLE-prone mice has no effect on apoptosis pathway genes in splenic immune cells.** Pro-/anti-apoptotic scores of each immune cell by pro- and anti- apoptosis gene signatures from Merino et al. (2013) within the $PRLR^+$ and $PRLR^-$ fractions. Data are from 4 *MRL-lpr* SLE-prone mice each in control SMO and LFPRLR SMO-treated groups.

***Fig S16.*** *Gating strategy for flow cytometry analysis of mature B cells*

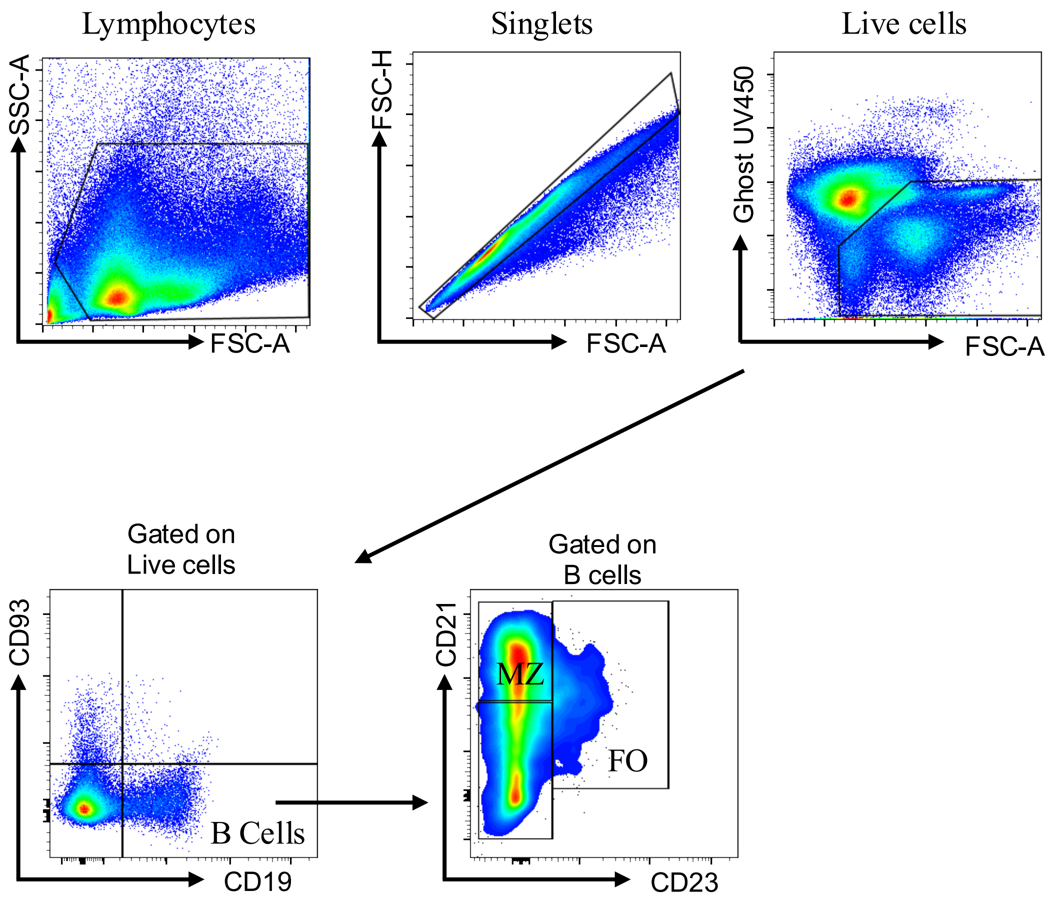


**Fig S16. Gating strategy for flow cytometry analysis of mature B cells.** From the lymphocyte cluster singlets were gated followed by selection of live cells (Ghost-UV450). B cells were identified as CD19$^{+}$ and further subdivided into transitional (CD93$^{+}$) and mature subsets. Mature B cells were classified as Marginal Zone (MZ; CD21$^{brigth}$CD23$^{low}$) and follicular (FO; CD21$^{int}$CD23$^{bright}$)

***Fig S17.*** *LFPRLR SMO reduces class-switched splenic B cells and alters homeostasis of CD11c/CD11b B-cell subsets including extrafollicular B cells*

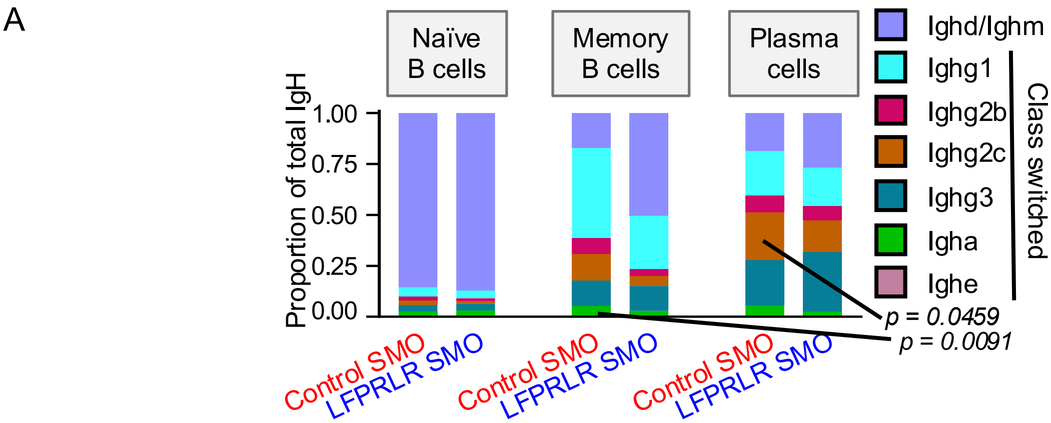


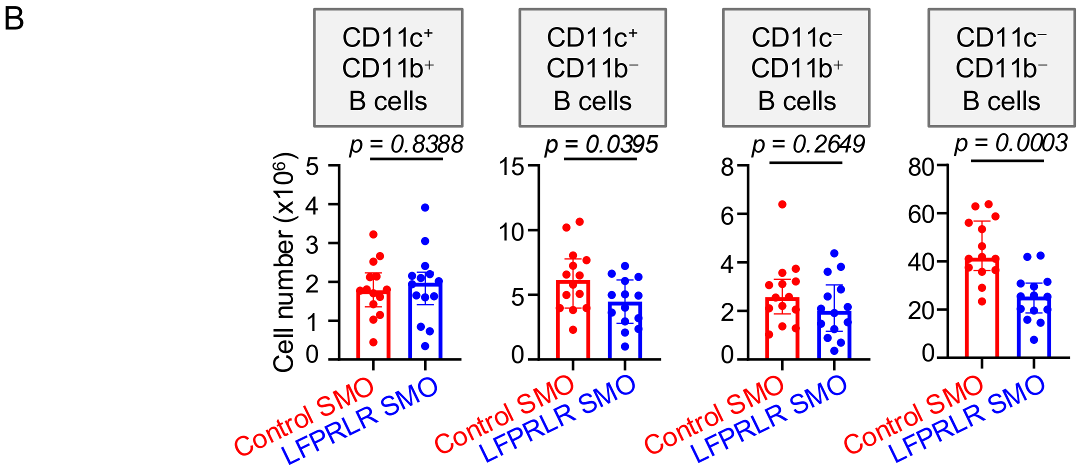


**Fig. S17. LFPRLR SMO reduces class-switched splenic B cells and alters homeostasis of CD11c/CD11b B-cell subsets including extrafollicular B cells.** (**A**) Relative proportions, calculated by taking the average of each mouse in each group, of class-switched (IgG/A/E) vs non-switched (IgD/M) naïve, memory, and plasma splenic B-cell subsets in control SMO (n=4) and LFPRLR SMO-treated (n=4) *MRL-lpr* SLE-prone mice by scRNA-seq. (**B**) Numbers of CD11c/CD11b B-cell subsets within the $CD19^{+}$ splenic B cell fraction by flow cytometry in control SMO (n=14) and LFPRLR SMO-treated (n=44) *MRL-lpr* SLE-prone mice. Data are represented as median ± interquartile range. p-values Mann-Whitney U test.

*Fig S18.* *Gating strategy for flow cytometry analysis of age-associated B cells*

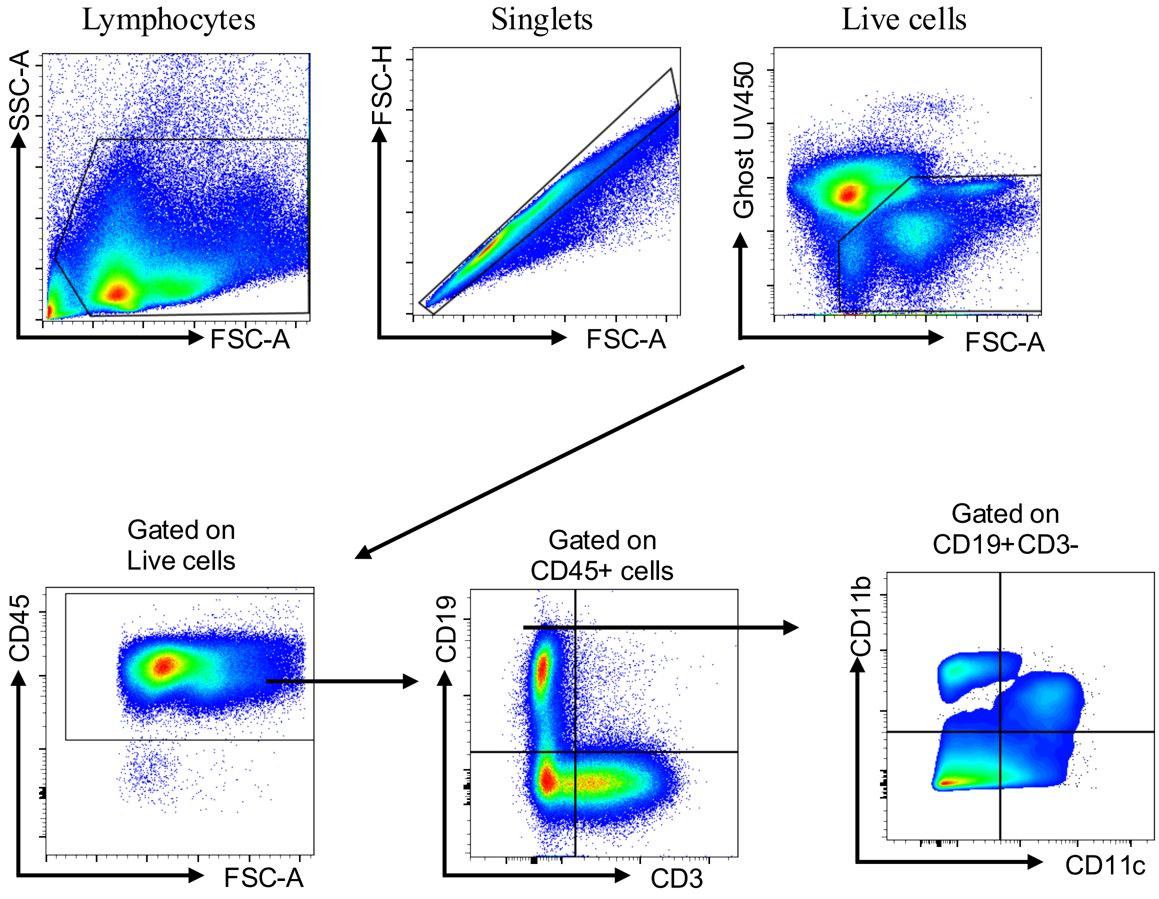


**Fig S18. Gating strategy for flow cytometry analysis of age-associated B cells.** From the lymphocyte cluster singlets were gated followed by selection of live cells (Ghost-UV450). B cells were identified as $CD19^+$ B cells and $CD3^+$ T cells, and further subdivided into $CD11c^+CD11b^-$, $CD11c^+CD11b^+$, $CD11c^-CD11b^+$, $CD11c^-CD11b^-$

***Fig S19.*** *Gene-expression profile of splenic B-cell subsets in healthy mice and SLE mice treated with control or LFPRLR SMO*

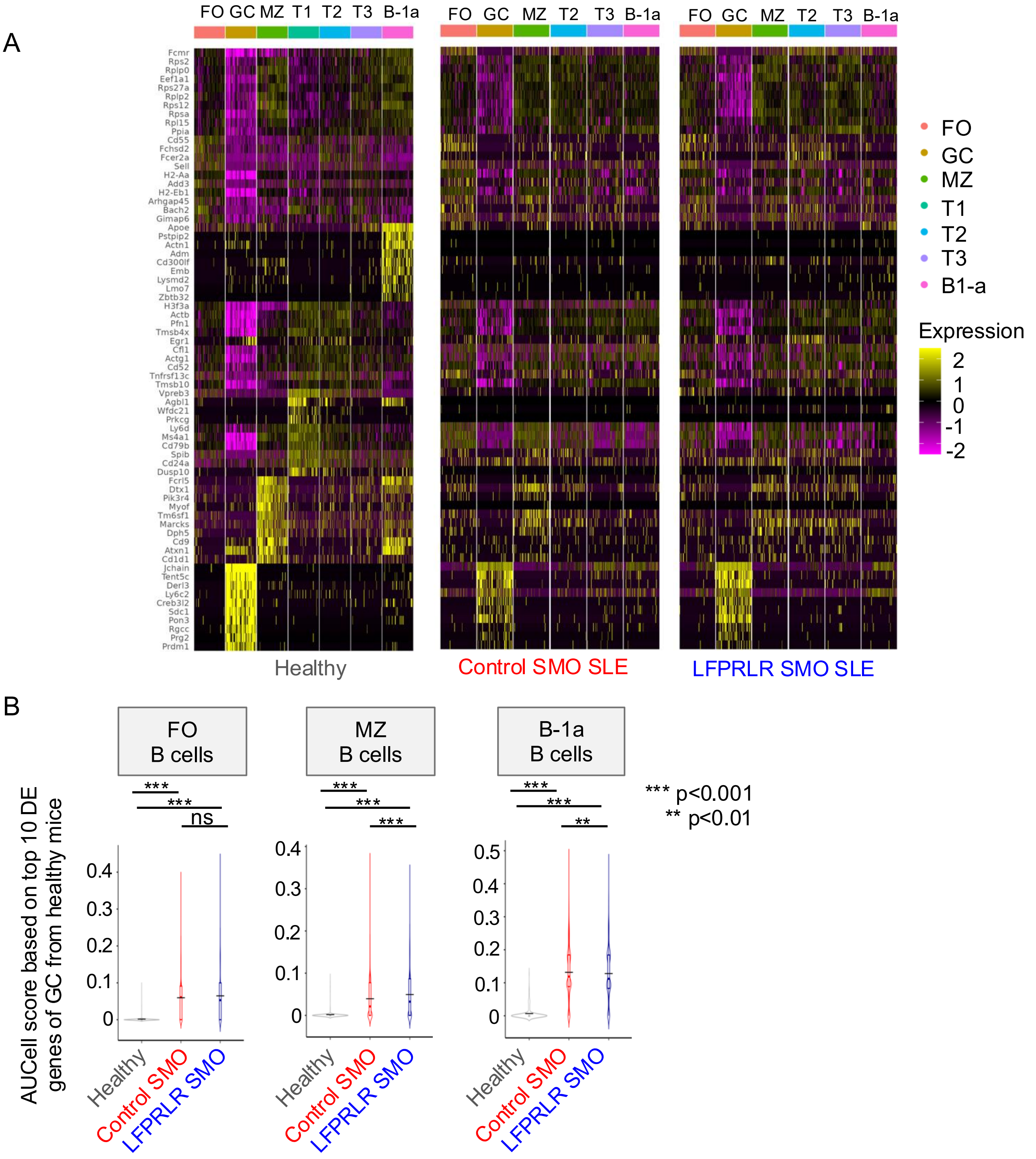


**Fig S19. Gene-expression profile of splenic B-cell subsets in healthy mice and SLE-prone mice treated with control or LFPRLR SMO.** (**A**) Heatmaps showing the expression of selected genes in splenic B-cell subsets from healthy (C57BL/6J, n=3), control SMO-treated (n=4) and LFPRLR SMO-treated (n=4) *MRL-lpr* SLE-prone mice. (**B**) AUCell score of FO, MZ, B-1a splenic B cell subsets in healthy mice (n=3), and control SMO- (n=4) and LFPRLR SMO- (n=4) treated MRL-lpr SLE-prone mice. AUCell score was calculated based on the top 10 differential expressed (DE) genes for each subset derived from healthy mice. Median ± interquartile range (black line depicts mean of each group), p-value: Mann-Whitney U test. FO, follicular B cells; GC, Germinal Center B cells; MZ, Marginal Zone B cells; T1, Transitional 1 B cells; T2, Transitional 2 B cells; T3, Transitional 3 B cells.

*Fig S20.* *LFPRLR knockdown for 8 weeks in SLE-prone mice has no effect on urinary protein and calciuria levels*

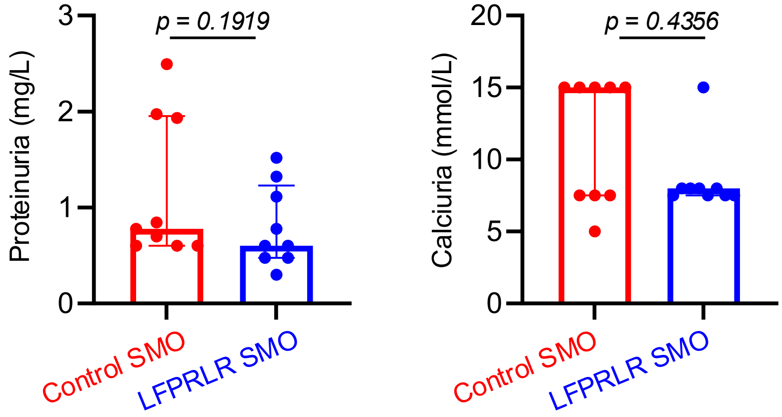


**Fig S20. LFPRLR knockdown for 8 weeks in SLE-prone mice has no effect on urinary protein and calciuria levels .** Proteinuria (mg/L) quantified by Bradford assay and Calciuria (mmol/L) assessed semi-quantitatively using multiparameter urine sticks in control SMO- (n=9) and LFPRLR SMO- (n=9) treated *MRL-lpr* mice. Mice were treated for 8 weeks and were 14 weeks old at the time of quantification of protein and calcium in the urine. Data are depicted as median ± interquartile range, p-values: Mann-Whitney U test.

**Table S1:** *List of SLE patient samples used in the study*

| Patient ID | Age | Sex | Tissue Type | Treatment duration | Current SLE treatment type | SLEDAI Score |
|---|---|---|---|---|---|---|
| 135502 | 41 | F | PBMC | > 2 months | Hydroxychloroquine | 6 |
| 95419 | 66 | F | PBMC | > 2 months | Hydroxychloroquine | 7 |
| 32573 | 63 | F | PBMC | > 2 months | Hydroxychloroquine | 6 |
| 132588 | 40 | F | PBMC | > 2 months | Methotrexate, Hydroxychloroquine | 0 |
| 23150 | 61 | F | PBMC | ≤ 2 months | Betamethasone, Hydroxychloroquine | 8 |
| 25093 | 53 | F | PBMC | ≤ 2 months | No current immunosuppressant or immunomodulator therapy | 6 |
| 33889 | 56 | F | PBMC | > 2 months | Prednisone | 10 |
| 100365 | 59 | F | PBMC | > 2 months | Prednisone, Celebrex, Leflunomide, Enbrel | 4 |
| 30045 | 69 | F | PBMC | > 2 months | Prednisone, Methotrexate | 6 |

SLEDAI score indicates disease activity: no activity (SLEDAI=0), mild activity (SLEDAI=1 to 5), moderate activity (SLEDAI=6 to 10), high activity (SLEDAI=11 to 19), and very high activity (SLEDAI≥20).

**Table S2:** *List of oligonucleotide sequences used in the study*

**a.** Splice modulating oligomer (SMO) (m= mouse, h=human)

| SMO | Sequence (5’→3’) |
|---|---|
| Control SMO (for mice and humans) | 5’-AGACGAGATTCGATCGGAGTA-3’ |
| mLFPRLR SMO | 5’-GCCCTTCTATTGAAACACAGATACA-3’ |
| hLFPRLR SMO | 5’-GCCCTTCTATTAAAAACACAGACACA-3’ |

**b. Quantitative RT-PCR**

| Primer | Sequence (5’→3’) |
|---|---|
| hUBB-F | 5’-GCCGCACTCTTTCTGACTACAAC-3’ |
| hUBB-R | 5’-ACCTCCAGAGTGATGGTCTTGC-3’ |
| hIFI35-F | 5’-CACGATCAACATGGAGGAGTGC-3’ |
| hIFI35-R | 5’-GGCAGGAAATCCAGTGACCAAC-3’ |
| hIFI44-F | 5’-GTGAGGTCTGTTTTCCAAGGGC-3’ |
| hIFI44-R | 5’-CGGCAGGTATTTGCCATCTTTCC-3’ |
| hIFITM3-F | 5’-CTGGGCTTCATAGCATTCGCCT-3’ |
| hIFITM3-R | 5’-AGATGTTCAGGCACTTGGCGGT-3’ |
| hIRF7-F | 5’-CCACGCTATACCATCTACCTGG-3’ |
| hIRF7-R | 5’-GCTGCTATCCAGGGAAGACACA-3’ |
| hMX1-F | 5’-GGCTGTTTACCAGACTCCGACA-3’ |
| hMX1-R | 5’-CACAAAGCCTGGCAGCTCTCTA-3’ |
| hMX2-F | 5’-AAAAGCAGCCCTGTGAGGCATG-3’ |
| hMX2-R | 5’-GTGATCTCCAGGCTGATGAGCT-3’ |
| hOAS1-F | 5’-AGGAAAGGTGCTTCCGAGGTAG-3’ |
| hOAS1-R | 5’-GGACTGAGGAAGACAACCAGGT-3’ |
| hOASL-F | 5’-GTGCCTGAAACAGGACTGTTGC-3’ |
| hOASL-R | 5’-CCTCTGCTCCACTGTCAAGTGG-3’ |
| hSTAT1-F | 5’-CCGTTTTCATGACCTCCTGT-3’ |
| hSTAT1-R | 5’-TGAATATTCCCCGACTGAGC-3’ |
| hSTAT2-F | 5’-CAGGTCACAGAGTTGCTACAGC-3’ |
| hSTAT2-R | 5’-CGGTGAACTTGCTGCCAGTCTT-3’ |
| hLFPRLR-F | 5’-TCCAGGTATGTGGGTTTCAT-3’ |
| hLFPRLR-R | 5’-GATTTGATGCTCATCTGTTGGA-3’ |
| hSF1aPRLR-F | 5’-TGGACTGTGGTCAATGTTGC-3’ |
| hSF1aPRLR-R | 5’-GATAGTGAGGACCAGCATCTAATG-3’ |

| | |
|---|---|
| hSF1bPRLR-F | 5’-CAACATCAAGGGGTCACCTC-3’ |
| hSF1bPRLR-R | 5’-CATGAATGATACAACCGTGTGG-3’ |

**Table S3:** *List of human flow cytometry antibodies used in the study*

| Marker | Fluorochrome | Clone | Source | Titer |
|---|---|---|---|---|
| CD197 (CCR7) | Spark NIR 685 | G043H7 | Biolegend | 1:20 |
| CD14 | Spark Blue 550 | 63D3 | Biolegend | 1:20 |
| CD185 (CXCR5) | BV480 | RF8B2 | BD Bioscience | 1:20 |
| CD27 | APC-R700 | M-T271 | BD Bioscience | 1:20 |
| CD138 | PE-Fire 640 | MI15 | Biolegend | 1:40 |
| CD38 | BUV737 | HIT-2 | BD Bioscience | 1:40 |
| CD19 | APC-H7 | HIB19 | BD Bioscience | 1:40 |
| CD11c | RB744 | B-ly6 | BD Bioscience | 1:40 |
| CD4 | BUV496 | SK3 | BD Bioscience | 1:40 |
| CD57 | eFluor 450 | TB01 | eBioscience | 1:40 |
| CD8 | BV510 | SK1 | Biolegend | 1:40 |
| Ki-67 | BUV805 | B56 | BD Bioscience | 1:80 |
| CD16 | RB613 | 3G8 | BD Bioscience | 1:80 |
| IgD | BB700 | IA6-2 | BD Bioscience | 1:80 |
| CD3 | APC-Fire 810 | SK7 | Biolegend | 1:160 |
| CD183 (CXCR3) | PE-Cy5 | G025H7 | Biolegend | 1:160 |
| CD123 | RB705 | 7G3 | BD Bioscience | 1:160 |
| CD56 | BUV615 | NCAM16.2 | BD Bioscience | 1:160 |
| CD137 | BV421 | 4B4-1 | BD Bioscience | 1:160 |
| CD196 (CCR6) | PE-Fire 700 | G034E3 | Biolegend | 1:160 |
| CD194 (CCR4) | Pe-Fire 810 | L291H4 | Biolegend | 1:160 |
| CD69 | BV650 | FN50 | Biolegend | 1:160 |
| Live/Dead UV blue | UV Blue | _ | Thermo Fisher Scientific | 1:500 |

**Table S4:** *List of mouse flow cytometry antibodies used in the study*

| Marker | Fluorochrome | Clone | Source | Titer |
|---|---|---|---|---|
| CD3 | BUV395 | 17A2 | BD Biosciences | 1:100 |
| CD4 | BV510 | RM4-5 | BioLegend | 1:100 |
| CD8 | BV711 | 53-6.7 | BioLegend | 1:100 |
| CD11b | FITC | M1/70 | BioLegend | 1:100 |
| CD11c | PE-Cy7 | N418 | BioLegend | 1:100 |
| CD19 | PerCP-Cy5.5 | 1D3 | BD Biosciences | 1:140 |
| CD19 | APC | 1D3 | BD Biosciences | 1:100 |
| CD21/35 | PE | 7E9 | BioLegend | 1:100 |
| CD23 | Biotin | B3B4 | BioLegend | 1:100 |
| CD27 | BV650 | LG.3A10 | BioLegend | 1:100 |
| CD45 | PerCP | 30-F11 | BioLegend | 1:100 |
| CD93 | APC | AA4.1 | BioLegend | 1:100 |
| B220 | APC-Cy7 | RA3-6B2 | BioLegend | 1:200 |
| Gr-1 | AF700 | RB6-8C5 | BioLegend | 1:100 |
| IgD | PE-Cy7 | 11-26c.2a | BioLegend | 1:100 |
| IgM | FITC | II/41 | Thermo Fisher Scientific | 1:100 |
| NK1.1 | BV605 | PK136 | BioLegend | 1:100 |
| NKp46 | PE | 29A1.4 | BD Biosciences | 1:100 |
| PDCA1 | BV750 | 927 | BD Biosciences | 1:100 |
| Streptavidin | BV711 | — | BD Biosciences | 1:100 |
| Ghost Dye | UV450 | — | Tonbo Biosciences | 1:100 |

**Table S5:** *List of human flow cytometry antibodies to confirm the generation of mature B-cell subsets from human PBMCs*

| Marker | Fluorochrome | Clone | Source | Titer |
|---|---|---|---|---|
| CD19 | BUV661 | HIB19 | BD Biosciences | 1:125 |
| CD27 | PE | M-T271 | BD Biosciences | 1:125 |
| CD38 | BV786 | HIT2 | BD Biosciences | 1:125 |
| CD3 | BUV805 | UCHT1 | BD Biosciences | 1:125 |
| CXCR4 | PECY5 | 12G5 | BioLegend | 1:125 |
| CD138 | BV510 | MI15 | BD Biosciences | 1:60 |
| IgD | BV650 | IA6-2 | BD Biosciences | 1:100 |
| CD269 (BCMA) | PECY7 | 19F2 | BioLegend | 1:125 |
| HLA-DR | BUV737 | G46-6 | BD Biosciences | 1:100 |
| IgG | BV421 | G18-145 | BD Biosciences | 1:125 |
| Ghost Dye | UV450 | — | Tonbo Biosciences | 1:100 |

**Table S6:** Software and packages used in the analysis

| Name | Version | URL |
|---|---|---|
| Cellranger | 7.2.0 | https://www.10xgenomics.com/support/software/cell-ranger/latest |
| samtools | 1.16.1 | https://www.htslib.org/ |
| STAR | 2.7.6a | https://github.com/alexdobin/STAR |
| Picard | 2.26.11 | https://broadinstitute.github.io/picard/ |
| HTSeq-count | 0.13.5 | https://htseq.readthedocs.io/en/release_0.11.1/count.html |
| DESeq2 | 1.42.1 | https://bioconductor.org/packages/release/bioc/html/DESeq2.html |
| DoubletFinder | 2.0.3 | https://github.com/chris-mcginnis-ucsf/DoubletFinder |
| Seurat | 5.2.1 | https://satijalab.org/seurat/ |
| clusterProfiler | 4.10.1 | https://bioconductor.org/packages/release/bioc/html/clusterProfiler.html |
| ComplexHeatmap | 2.18.0 | https://bioconductor.org/packages/release/bioc/html/ComplexHeatmap.html |
| scMRMA | 1.0.0 | https://github.com/JiaLiVUMC/scMRMA |
| SingleR | 2.10.0 | https://www.bioconductor.org/packages/release/bioc/html/SingleR.html |
